# Free-water volume fraction increases non-linearly with age in the white matter of the healthy human brain

**DOI:** 10.1101/2022.10.06.510800

**Authors:** Tomasz Pieciak, Guillem París, Dani Beck, Ivan I. Maximov, Antonio Tristán-Vega, Rodrigo de Luis-García, Lars T. Westlye, Santiago Aja-Fernández

## Abstract

The term free-water volume fraction (FWVF) refers to the cerebrospinal and interstitial fluids in the extracellular space of the white matter (WM) of the brain, which has been demonstrated as a sensitive biomarker that correlates with the cognitive performance and the neuropathological processes modifying the interstitial extracellular spaces. It can be quantified by properly fitting the isotropic compartment of the magnetic resonance (MR) signal in diffusion-sensitized sequences. Using *N* = 287 healthy subjects aged 25-94, this study examines in detail the evolution of the FWVF in the human brain WM across the adult lifespan, which has been previously reported to exhibit a positive trend. We found evidence of a noticeably non-linear gain after the sixth decade of life, with a region-specific variate and varying change rate of the FWVF parameter with age, at the same time a heteroskedastic pattern across the adult lifespan is suggested. On the other hand, the FW-compensated MR signal leads to a region-dependent flattened age-related evolution of the mean diffusivity (MD) and fractional anisotropy (FA), along with a considerable reduction in their variability, as compared to standard studies conducted over the raw MR signal. This way, our study provides a new perspective on the trajectory-based assessment of the brain and explains the source of the variations observed in FA and MD parameters across the lifespan with previous studies with the standard diffusion tensor imaging.

## Introduction

Brain imaging has provided strong evidence that the human brain varies throughout development and aging (Sowel et al., 2003; Shaw et al., 2008; Tamnes et al., 2010; Westlye et al., 2010; Lebel et al., 2012; Lebel and Deoni, 2018; Bethlehem et al., 2022; Grotheer et al., 2022). Although the neurobiological underpinnings are manifold, including axon myelination and synaptogenesis (Huttenlocher et al., 1997; Nave, 2010), they are especially noticeable in the early years of life (Giedd et al., 1999; Lebel and Deoni, 2018; Bethlehem et al., 2022). A recent multiple-sites study (Bethlehem et al., 2022) reported that, after childhood, there is a systematic reduction in gray matter and white matter (GM, WM) volumes, coupled with ventricles augmentation and an associated apparent thinning of the cerebral cortex. In addition, it has been found that differences in brain properties also exist between male and female brains, accentuating with age (Sowell et al., 2007; Bethlehem et al., 2022). Accurate delineations of age-related trajectories of brain structural features can help us better understand its functioning and discern which patterns can be considered as “normative trajectories” (Bethlehem et al., 2022; Rutherford et al., 2022). Deviations from the regular patterns could be indicators of neurodevelopmental disorders as well as relevant markers for the diagnosis and prognosis of brain pathologies (Planche et al., 2022).

Although age-related structural differences in the human brain were initially investigated in *post-mortem* cross-sectional studies (Dekaban et al., 1978), more recent research has observed these variations *in vivo* with magnetic resonance imaging (MRI) (Sowel et al., 2003; Gogtay et al., 2004; Bethlehem et al., 2022). The diversity of MRI-derived modalities has enabled us to inspect the properties of various tissues and brain regions across the lifespan from both structural and microstructural perspectives (MacDonald and Pike, 2021). As previously stated, the WM is one of the major brain tissue classes, and it effectively changes with age, partly due to myelin sheath formation around neuronal axons during development and early childhood, but also due to axonal shrinkage with the progressive oligodendrocyte death, demyelination with the formation of myelin spheroids and myelin debris accumulation during senescence (Peters 2002; Nave, 2010; Hill et al., 2018; Lebel and Deoni, 2018). The fluctuations in microstructural properties of the brain are associated with a range of physical and inherent biological processes, including cognition (Maillard et al., 2019), normal aging (Westlye et al., 2010; Lebel et al., 2012; Cox et al., 2016; Beck et al., 2021; Kiely et al., 2022) and the development of neurodegenerative diseases (Kubicki et al., 2007; Raghavan et al., 2021).

Diffusion magnetic resonance imaging (dMRI) has been widely applied to the study of healthy brain development and the aging (Bennett et al., 2010; Burzynska et al., 2010; Tamnes et al., 2010; Westlye et al., 2010; Lebel et al., 2012; Cox et al., 2016; Beck et al., 2021; Grotheer et al., 2022), as well as neurological disorders that occur throughout life (Head et al., 2004; Cetin-Karayumak et al., 2020). dMRI allows the characterization of the diffusivity of water molecules within tissues, providing information about the microscopic configuration and structural connectivity of the brain. The most relevant feature of dMRI is its ability to measure the directional variance of water diffusivity in a single voxel, i.e. diffusion anisotropy (Basser et al., 1994). Different mathematical representations are used to describe the relationship between the acquired dMRI signal and the properties of the tissue under study. The most common way to estimate the diffusivity properties is via the single-component diffusion tensor imaging (DTI), representing the restricted diffusion that follows a Gaussian-like behavior (Basser et al., 1994; Westin et al., 2002) and the derived scalars, such as the fractional anisotropy (FA) or the mean diffusivity (MD). These single-component DTI-based measures exhibit a well-documented curvilinear U-form across the lifespan in the healthy human brain, i.e. a rapid growth of FA parameter in childhood and adolescence and a systematic decline after the peak value, consistent with the MD measure, which reveals the opposite trend (Westlye et al., 2010; Lebel et al., 2012; Beck et al., 2021).

Alternatively, there are other dMRI signal representations that can be used to quantify diffusion properties, with particular focus on the WM (Jensen et al., 2005; Özarslan et al., 2013; Tristán-Vega and Aja-Fernández, 2021). Despite their far-reaching abilities to represent Gaussian or non-Gaussian diffusion, the simplest extension of DTI is to assume that, inside the brain tissues, we can consider two different components, one related to the neural cells representing the restricted diffusion (the part that can be measured by the DTI) and another presenting the free-water diffusion, i.e., the diffusion that is neither restricted nor hindered (Pierpaoli and Jones, 2004; Pasternak et al., 2009; Rydhög et al., 2017). An attractive characteristic of such a representation is the free-water volume fraction (FWVF) parameter, which is defined as the part of free diffusion unveiled as the cerebrospinal fluid (CSF) and interstitial fluid in the extracellular space of WM and GM from the total amount of diffusion (Pasternak et al., 2012; Rydhög et al., 2017).

Several recent studies have shown that the variations in the FWVF may depend on cognitive performance (Maillard et al., 2019), or occur due to neuroinflammation or atrophy of the aging brain (Metzler-Baddeley et al., 2012; Pasternak et al., 2012; Kubicki et al., 2019). Changes in the FWVF may also reflect the response to pathological processes that modify the interstitial extracellular spaces (Chad et al., 2018). Examples of conditions leading to increased FWVF values include parenchymal edema, such as those around tumors (Pasternak et al., 2009), Parkinson’s disease (Ofori et al., 2015), first-episode of psychosis (Lyall et al., 2018), Alzheimer’s disease (Bergamino et al., 2021) and schizophrenia (Carreira Figueiredo et al., 2022). The main implication of recovering FWVF information is related to the palliation of partial volume effects created by free-water diffusion (Metzler-Baddeley et al., 2012), given that DTI-based metrics are often contaminated with such, and its elimination results in the reduction in the number of false-positive streamlines in DTI-based tractography (Pasternak et al., 2009) and an improvement of the metrics’ reproducibility (Albi et al., 2017). In addition, recent studies have suggested that the FWVF parameter is positively correlated with age in healthy humans’ (Chad et al., 2018) and adult rhesus monkeys’ (Kubicki et al., 2019) brains WM, and could therefore affect the evolution in DTI-related parameters across the lifespan (Metzler-Baddeley et al., 2012; Chad et al., 2018; Kubicki et al., 2019). All in all, however, a detailed description of the evolution of FWVF across the adult lifespan, and its possible effects on diffusional properties of the brain WM have not been clarified.

The FWVF can be estimated from dMRI data sets in a number of different ways. Biophysical models (Cox et al., 2016; Raghavan et al., 2021) can imitate the underlying geometry of intra-voxel neural tissue, including the extracellular water diffusion with reasonable accuracy, but they usually require dedicated dMRI data with specific acquisition protocols, which are not likely available in general-purpose databases. On the other hand, the bi-tensor scheme based on the aforementioned two-component representation (Pierpaoli and Jones, 2004; Pasternak et al., 2009) can resolve the FWVF even from standard dMRI acquisitions in case proper numerical optimization schemes are set up (Pasternak et al., 2009; Pasternak et al., 2012; Golub et al., 2020). This approach, however, suffers from a positive bias observed in the voxels presenting complex architectures such as fiber crossings (Tristán-Vega et al., 2022), which are known to be present in up to 90% of the white matter voxels (Jeurissen et al., 2013). Alternatively, we will use the so-called spherical means technique (Kaden et al., 2016; Tristán-Vega and Aja-Fernández, 2021), in which the signals obtained for all orientations of the diffusion gradients with the same magnitude are averaged together. This way, the impact of a varying number of crossing fibers and confounding factors (i.e., the number of diffusion-sensitizing gradients) is removed, and the FWVF can be unbiasedly estimated over the WM whenever two different gradient magnitudes are included in the protocol (Tristán-Vega et al., 2022).

This paper investigates how the FWVF varies over the adult lifespan in the WM of the healthy human brain using cross-sectional and longitudinal multi-shell dMRI data. The main contributions of this work are the following: we demonstrate that (1) the FWVF varies non-linearly in the healthy human brain WM across the adult lifespan with a significant positive gain after the sixth decade of life; (2) the FWVF exhibits a heteroskedastic pattern with age; (3) we observe a region-specific dependence in the FWVF evolution, with the most pronounced shift in the cingulum and the genu of the corpus callosum and (4) the presence of a spatial gradient of the water fraction. We observe that (5) the FWVF displays correlations with the brain, cerebral WM and ventricles volumes highlighting the heteroskedastic nature of the parameter as a function of the volume-based feature. Finally, (6) we have detected a huge influence of the FWVF on the variabilities of measures derived from DTI such as MD and FA across the adult lifespan. This variability can be significantly reduced when removing the bias introduced by the free water.

## Methods

### Summary in modeling the free-water, structural and diffusional properties of the brain over the population study

The sample population used in the study includes 287 healthy subjects (178F/109M) and has been divided into eight groups, each aggregating the subjects within a ten-year interval (see Table 1). For a single subject, we calculated: (1) the volume-based features of the brain from T1-weighted scans using the FreeSurfer software suite, (2) the FWVF parameter *f* from two-shell dMRI data acquired at *b* = {1000, ^2^000} s/mm^2^ using the spherical means approach (Tristán-Vega et al., 2022), (3) the standard DTI-based MD and FA measures from single-shell dMRI data at *b* = 1000 s/mm^2^ and (4) the FW corrected equivalent DTI-based measures from single-shell dMRI data at *b* = 1000 s/mm^2^ given the estimated FWVF *f* in the two-component representation (Eq. (3)). Twenty three regions of interest (ROIs) from the Johns Hopkins University (JHU) WM atlas (Mori et al., 2005) are used in the study in order to evaluate the region-specific properties of FWVF and DTI-based measures of the brain. In Fig. 1b and Supplementary Fig. 1a,c, the ROIs considered in the study are drawn in the standard space over the selected coronal and axial slices of the T1-weighted template, and additionally visualized through 3D models. The region-specific age-related trajectories in FWVF and DTI-based parameters across the adult lifespan are modeled using the quantile regression (QR) technique (Koenker and Hallock, 2001). The QR models the conditional median of the parameter *f* given the explanatory variable *Age* (Model 1, Eq. (4)) being the subject’s age at the scan time and optionally *FSIQ* (Full Scale Intelligence Quotient) (Model 2, Eq. (5)) or *Sex* (a binary variable) (Model 3, Eq. (6)). Similarly, we model the variations in the standard DTI and FW corrected DTI measures (i.e., FA, MD) using the second-order QR (Model 1) given the explanatory variable *Age*. The first-order QR (Model 4, Eq. (7)) is employed to model the variations in the FWVF parameter *f* as a function of volume-based features, i.e., the brain volume, cerebral WM volume and ventricles volume.

**Table 1.**
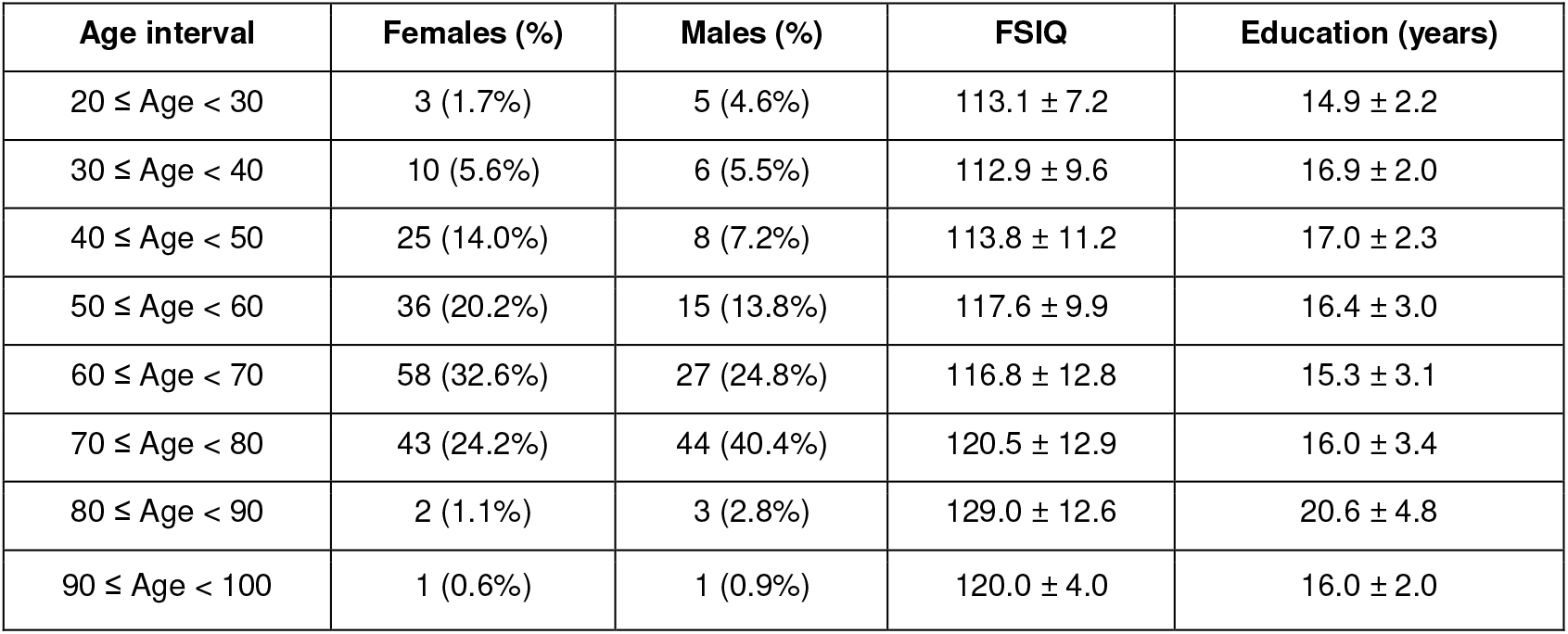
Population distribution used in the study (*N* = 287 healthy cases, 178F/109M) including the FSIQ and years of education (mean ± std. dev).

**Figure 1.**
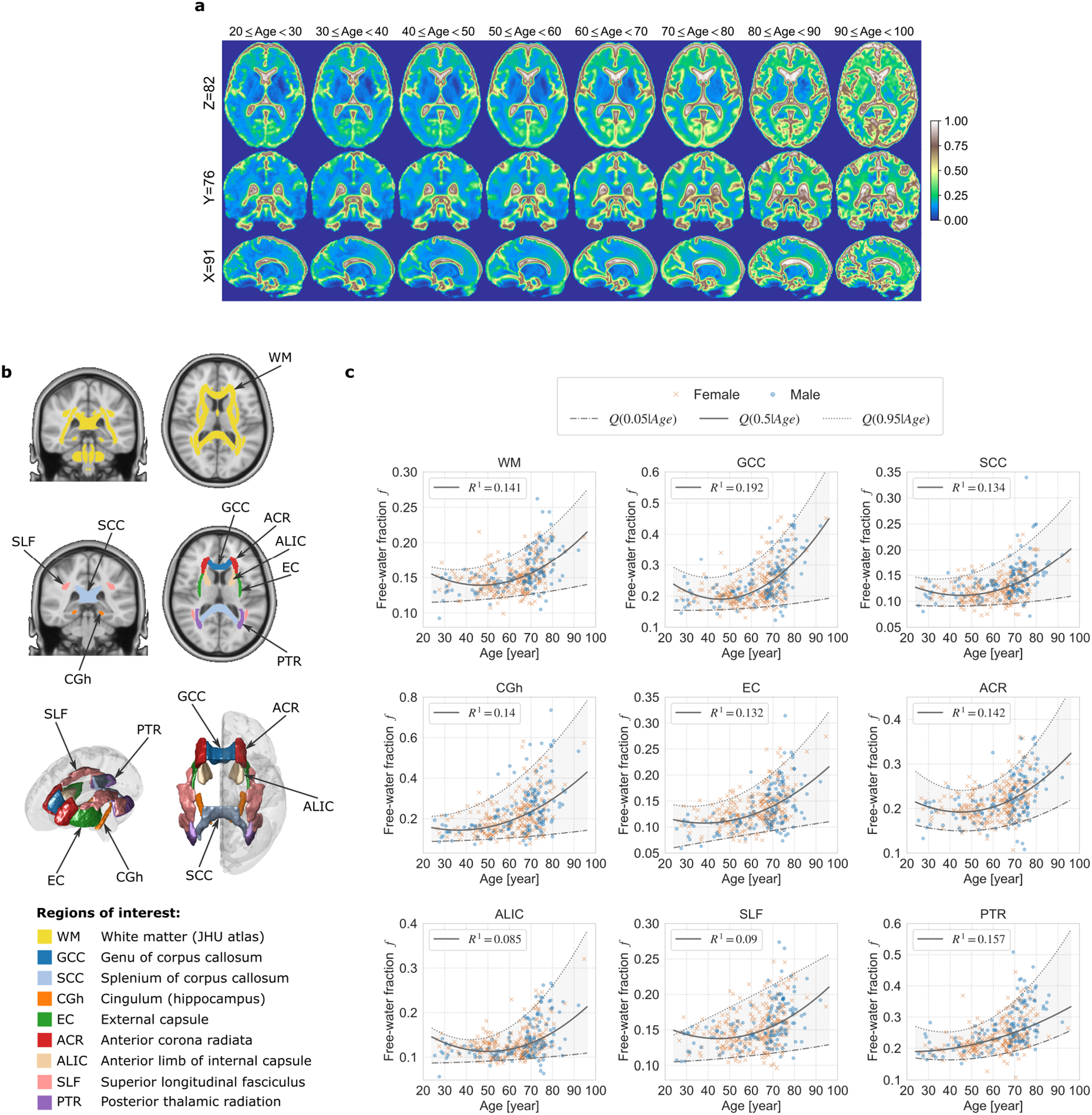
The trajectories of the FWVF parameter across the study age span shown over different regions of interest (ROIs). **a** Visual inspection of the FWVF *f* across age intervals presented separately in axial (Z=82), coronal (Y=76) and sagittal (X=91) planes. The images are displayed in the standard space after non-linear warpings computed with the single-component FA registration from the subjects’ native spaces. Each figure presents a median image of the FWVF *f* computed voxel-wise across all subjects available in a specified range (only cross-sectional samples are used for this experiment). **b** The ROIs used in the study were retrieved from the JHU DTI-based atlas (Mori et al., 2005) and visualized in the standard space over the T1-weighted coronal and axial slices, and using 3D models. **c** Estimated FWVFs and their trajectories computed with a quantile regression technique. An individual marker delivers a median value of the FWVF *f* computed from the ROI in the native space of a single subject. The experiment uses cross-sectional and longitudinal samples. The quantile function *Q*(*τ*|*Age*) for a single ROI is shown under three quantiles, i.e., the solid line presents *Q*(*τ*|*Age*) under *τ* = 0.5 (50th percentile), the lower thin dashed-dotted line indicates *Q*(*τ*|*Age*) under *τ* = 0.05 (5th percentile), and the upper thin dotted line presents *Q*(*τ*|*Age*) under *τ*= 0.95 (95th percentile). All three were computed from median FWVFs. The regions between *Q*(0.05|*Age*) and *Q*(0.95|*Age*) were shaded for visualization purposes. The goodness-of-fit *R*^1^ at *τ* = 0.5 was computed for each region using the procedure introduced by Koenker and Machado (1999).

### Recruitment procedure, exclusion criteria and sample population

In total, *N* = 287 healthy subjects (178F/109M), aged (F/M mean ± std.dev: 60.6 ± 12.7/63.7 ± 15.0) were scanned after obtaining informed consent. Out of the 287 subjects, 99 were scanned again after periods between 10 and 30 months. The study was conducted in line with the Declaration of Helsinki and has been reviewed and approved by the Regional Committee for Medical and Health Research Ethics (REC), South East division in Norway (2014/694, 2009/2485). Inclusion criteria for the study included being above the age of 18 years and absence of alcohol and drug dependence. The exclusion criteria included neurological and mental disorders, and suffering from a previous head trauma. Participants were also excluded if they were not compatible for MRI scanning, including being pregnant at the time of data collection or having pacemakers or metal implants. The participants completed comprehensive clinical and neuropsychological/ cognitive assessments, blood sampling, and multimodal neuroimaging. The population was made up of largely ethnic Scandinavian and Northern European individuals. The demography across the age intervals including the FSIQ score and years of education are summarized in Table 1, while in Fig. 5, we visualize the population distribution of the study and time periods between two scans for 99 subjects.

### MRI data acquisition

All subjects were scanned at Oslo University Hospital with a 3T General Electric Discovery MR750 scanner (GE, Waukesha, WI) equipped with a 32 channel head coil and a maximum gradient strength of 40 mT/m.

#### Diffusion MRI

The multi-shell diffusion MRI data were collected using an echo-planar imaging (EPI) sequence with the following parameters: time echo TE=83.1 ms, repetition time TR=8150 ms, flip angle 90º, voxel size 2 × 2 × 2 mm^3^, field-of-view FOV 256 × 256 mm^2^, acquisition matrix 128 × 128 and 66 slices covering the brain. The data were acquired under two b-shells at *b* = 1000 s/mm^2^ and *b* = 2000 s/mm^2^ along with non-collinear 60 and 30 uniformly distributed diffusion gradient directions per shell, respectively. Each session included ten non-diffusion-weighted volumes (i.e., *b* = 0) and additionally seven non-diffusion-weighted volumes acquired in a reversed phase-encoding direction used to correct the susceptibility distortions. Total scan time was 8 min 58 sec.

#### T1-weighted MRI

The structural T1-weighted data were collected using a 3D inversion recovery prepared fast spoiled gradient recalled sequence (IR-FSPGR; BRAVO) using the following parameters: TE=3.18 ms, TR=8.16 ms, flip angle 12º, voxel size 1 × 1 × 1 mm^3^, FOV 256 × 256 mm^2^, acquisition matrix 128 × 128 and 66 slices covering the brain. Total scan time was 4 min 43 sec.

### Data quality assurance (QA) protocol

The initial population sample underwent a QA protocol, including testing for head motions, noise, and other acquisition-related artifacts. Additionally, the data sets were manually inspected if the temporal signal-to-noise-ratio parameter (Roalf et al., 2016) deviated ±2.5 standard deviation from the mean value.

### Data preprocessing

#### Diffusion MRI

The data were preprocessed using the pipeline adapted from Maximov et al. (2019), including noise removal using the Marčenko-Pastur principal component analysis technique (Veraart et al., 2016), Gibbs ringing artifacts correction, susceptibility-induced distortions, head movements and eddy current distortions correction using the FSL `eddy` tool. Eventually, the isotropic smoothing with the FSL `fslmath` was applied with a Gaussian kernel size of 1 × 1 × 1 mm^3^ to clean potential distortions after eddy current corrections.

#### T1-weighted MRI

The data were processed using the FreeSurfer 5.3, including non-uniformity intensity correction, computation the Talairach transform, skull stripping and the non-linear volumetric registration. The brain labeling was performed using the automatic subcortical segmentation using the probabilistic atlas (Fischl et al., 2002). The following volume-based features are considered in the study: total intracranial volume (TIV), brain volume, brain volume without ventricles and cerebral WM volume.

### Free-water volume fraction (FWVF) estimation

The FWVF parameter *f* was estimated from a two-shell diffusion MRI acquisition using the spherical means approach (Tristán-Vega et al., 2022) that assumes the diffusion is modeled using a continuous mixture of axis-symmetric second-order tensors in the space of orientations. The normalized dMRI signal is represented as follows (Tristán-Vega and Aja-Fernández, 2021; Tristán-Vega et al., 2022):

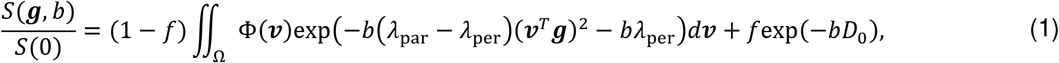

where *f* ∈ [0, 1] is the FWVF parameter, ***g*** is the diffusion sensitizing gradient direction, *b* is the *b*-value, Φ(***ν***) ≥ 0 is the convolution kernel, Ω = {***ν*** ∈ ℝ^3^: ∥***ν***∥ = 1}, *λ*_par_ and *λ*_per_ are the longitudinal and perpendicular diffusivities that satisfy the formula 0 ≤ *λ*_per_ ≤ *λ*_par_ ≤ *D*_0_ with *D*_0_ = 3.0 · 10^−3^ mm^2^/s being the apparent diffusion coefficient of free-water given for a water temperature of 37°*C* (Pasternak et al., 2009). The FWVF *f* from Eq. (1) was found by optimizing the following cost function:

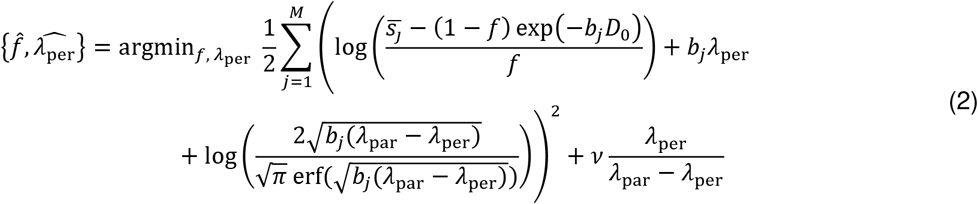

with 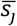 being the orientationally-averaged dMRI signal for *j*-th shell (*j* = 1, …, *M*) via the spherical harmonics decomposition at the order of *L* = 6 using an inverse linear problem with a Laplace-Beltrami regularization at *λ*= 0.001, *b*_*j*_ is *j*-th b-value (*b* ∈ {1000, 2000} s/mm^2^) and *v* = 0.06 is the penalty term that promotes the prolate convolution kernels. The Eq. (2) was solved using a constrained optimization via the gradient-projection method by fixing the parallel diffusion *λ*_par_ = 2.1 · 10^−3^ mm^2^/s.

### Diffusion tensor imaging (DTI) under a two-component representation

The DTI-based metrics were estimated under two variants, namely a standard approach proposed by (Basser et al., 1994) and a two-component representation illustrating the restricted diffusion via a diffusion tensor and free diffusion by a mono-exponential decay (Pierpaoli and Jones, 2004; Pasternak et al., 2009):

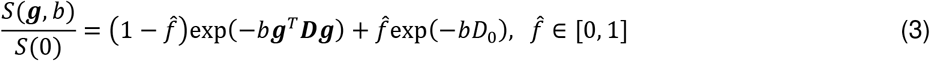

with 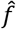 being the pre-estimated FWVF parameter using the spherical means approach (Tristán-Vega et al., 2022), and ***D*** is a symmetric semi-positive matrix of size 3 × 3 representing the restricted diffusion. The two-component representation given by Eq. (3) reduces to the standard DTI for 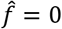.

To estimate the diffusion tensor ***D*** in either representation, we used the least squares approach solving the system of linear equations ***D*** = (***X***^*T*^***X***)^−1^***X***^*T*^ ***S*** with ***S*** being the vector of dMRI signals acquired at *b* = 1000 s/mm^2^, i.e., 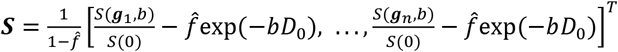 and ***X*** is the encoding gradient design matrix. Two measures were obtained from either representation, namely the MD and FA.

### Data registration and regions of interest retrieval

The FA parameters estimated using the standard DTI at *b* = 1000 s/mm^2^ were used to register the data to the standard space and retrieve the JHU WM atlas labels in the native subjects’ spaces (Mori et al., 2005). First, the FA parameters were affinely registered to the FSL template FMRIB58_FA via the FSL `flirt` tool (Analysis Group, FMRIB, Oxford, UK) using the normalized correlation cost function and a trilinear interpolation. We applied then a non-linear transformation (`fnirt`; FSL) and obtained a reverse warping to the subjects’ standard spaces via the FSL `invwarp`. The JHU WM regions of interest (Mori et al., 2005) were transformed to the subjects’ native spaces using a nearest-neighbor interpolation. In order to eliminate potential misregistration outliers due to a partial volume effect the regions of interest were shrunk with a morphological binary erosion operator with a cubic kernel of size 2 × 2 × 2.

### Statistical analyses

#### Quantile regression (QR)

The QR technique (Koenker and Hallock, 2001) was used to model the changes in the FWVF *f* and DTI-based parameters across the adult lifespan, and the relation between the DTI-based measures and the FWVF *f*. The QR estimates the conditional median of the parameter and allows for modeling its heteroskedastic nature without any theoretical assumptions on the distributional properties of the data.

To model the changes of the parameter (e.g., *f*, MD, FA) given the explanatory variable *Age* being the subject’s age at the scan time, and optionally to handle the continuous full-scale intelligence quotient variable (*FSIQ*) or the binary variable *Sex*, we used the following models:

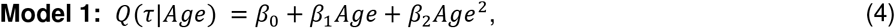

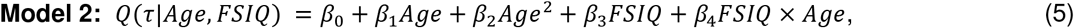

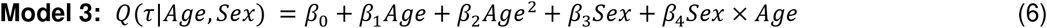

with *β*_*k*_ ∈ ℝ being the unknown parameters to be estimated. The models given by Eqs. (5) and (6) enable us to additionally include two-way interactions between the variables *Age* and *FSIQ*/*Sex*. The optimal order of the polynomial as a function of the variable *Age* (i.e., the first-order or the second-order) for each ROI was selected using the Akaike information criterion (AIC) (Akaike, 1974).

To analyze the changes of the FWVF *f* as a function of volume-based features, we used the first-order QR given by the formula:

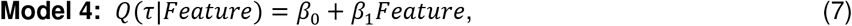

where *Feature ∈* {*Brain volume, Cerebral WM volu*m*e, Ventricles volume*} is the explanatory variable.

The model parameters ***β*** = [*β*_0_, …, *β*_*k*−1_] in Eqs. (4-7) under the quantile *r* were found using the following formula:

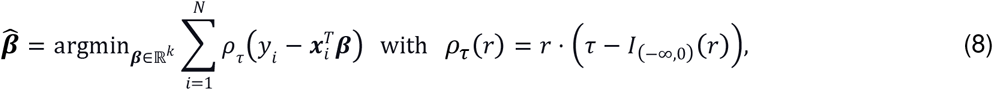

where ***x***_***i***_ = [1, *Age*_*i*_, …] is the *k*-th dimensional vector of covariates, y_*i*_ is the response variable from *i*-th subject of the study and *l*_(−*∞*,0)_(*r*) is the indicator function. The Eq. (8) is optimized using Brent’s method (Brent, 1973). For all models, we calculated the quantile function *Q*(*τ*|*Age*) under three quantiles *τ*, i.e., *τ* = 0.05 (5th percentile), *τ* = 0.5 (median), and *τ* = 0.95 (95th percentile).

#### Goodness-of-fit in QR

To calculate the goodness-of-fit of the model fitted with the QR framework under the quantile *τ*, we used the coefficient *R*^1^ introduced by Koenker and Machado (1999):

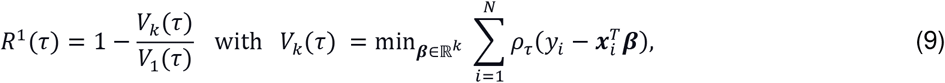

where *V*_1_(*τ*) and *V*_*k*_((*τ*) are the parameters calculated with the non-conditional model (i.e., intercept only) and the conditional model, respectively.

#### Doane’s formula

The number of bins *N*_*h*_ used to compute a single density plot given the sample of size *h* was calculated upon Doane’s formula (Doane, 1976):

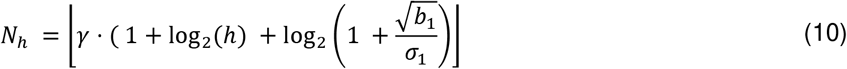

with the statistics 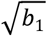 being the third standardized sample moment and 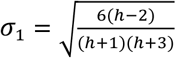 being the standard deviation of 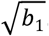. The scalar *γ* > 0 is the scaling factor and ⌊y⌋ is the function that provides the largest integer number less than or equal to *y*.

#### Breusch-Pagan (BP) test

The BP <u>statistical</u> test was used to assess the heteroskedasticity in the FWVF *f* as a function of volume-based parameters. The null hypothesis H_0_ of the test states that the error variances of the linear model are equal (i.e., the homoskedasticity present in the FWVF *f*), while the alternative hypothesis H_1_ states that the error variances are not equal, i.e., heteroskedasticity exists in the FWVF parameter (Breusch and Pagan, 1979). We used the studentizing correction to the BP test as suggested by Koenker (1981).

### Software

Diffusion MRI data were preprocessed with the FSL FMRIB Software Library v6.0 (Analysis Group, FMRIB, Oxford, UK.; https://fsl.fmrib.ox.ac.uk/fsl/fslwiki). The volumetric segmentation from T1-weighted data was carried out with the FreeSurfer 5.3 (recon-all command) image analysis suite (http://surfer.nmr.mgh.harvard.edu/). The free-water volume fraction was estimated with the dMRI-Lab toolbox (https://www.lpi.tel.uva.es/dmrilab) implemented in MATLAB (The MathWorks, Inc., Natick, MA), while the DTI-based parameters were estimated with home-delivered software written in Python 3.8.5 (https://www.python.org/) and NumPy 1.19.2 library (https://numpy.org/). All statistical analyses were carried out with GNU R 4.1.2 using the packages quantreg, DescTools and lmtest (https://cran.r-project.org/web/packages/) and statsmodels 0.12.0 (https://www.statsmodels.org/).

## Results

### Free-water volume fraction shows a non-linear trend with age

In this first section, we study the region-specific changes in the FWVF across adulthood and evaluate the possible effects from sex and intelligence variables. For the sake of illustration, a preliminary visual inspection of the FWVF parameter is performed for three different slices and for eight different age intervals across the adult lifespan (see Fig. 1a). Each map shows voxel-wise median aggregated water fractions in the common space from all subjects available in a specified range (see Table 1). We observe a non-uniform FWVF with higher values in the anterior part of the brain compared to the medial regions. These variations are especially pronounced beginning from the interval 60 ≤ *Age* < 70 (Fig. 1a, first row). Furthermore, we observe heightened FWVFs in the regions close to the CSF, highlighted with succeeding age intervals.

In Fig. 1c we illustrate the trajectories of the FWVF *f* over the whole WM area and eight selected regions of interest (ROIs) across the adult lifespan (see Supplementary Fig. 1 for the remaining 14 ROIs). Each marker relates to a single subject and displays a median value of *f* calculated from all samples available per ROI. The trajectories are modeled with the quantile regression (QR) technique (Koenker and Hallock, 2001) via a polynomial of the variable *Age* (see Model 1 in Methods). We present the quantile function *Q*(*τ*|*Age*) under three quantiles, i.e. *τ* = {0.05, 0.5, 0.95}. Further details, including the coefficients of the models for the different ROIs, their standard errors and *p*-values, are gathered in Supplementary Table 1.

In the whole WM area, we observe a non-linear behavior of the FWVF *f* across the adult lifespan that follows the model obtained at *τ* = 0.5 (50th percentile) given by the equation:

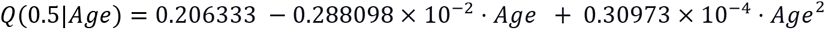

with *p* < .001 for all coefficients and the goodness-of-fit equals *R*^1^ = 0.141. The standard errors for the coefficients are 0.18^2^75 × 10^−1^ (*β*_0_), 0.58608 × 10^−3^ (*β*_1_) and 0.4711 × 10^−5^ (*β*^2^); note all three are one order of magnitude smaller than the corresponding coefficients. The age peak indicates the trend reversal in the FWVF over the WM at the age of 46y 6m. Regarding individual ROIs, we generally recognize a positive non-linear U-shaped trend represented by the second-order polynomial within most WM regions except for the ICP, where the first-order model slightly better represents the FWVF behavior in terms of the Akaike information criterion (AIC) (Akaike, 1974). The significance of the quadratic term in the second-order models varies across the regions within the range *p* < .001 and *p* < .05 excluding four regions, namely the PTR (*p* = .079), SCP (*p* = .^2^10), CT (*p* = .076), and the ML (*p* = .247). The trend reversal also varies in a region-specific manner spreading from 20y 7m (PTR) to 58y 4m (CT). Importantly, the QR technique allows for covering different quantiles over the data. We observe that the quantile functions *Q*(0.05|*Age*) and *Q*(0.95|*Age*) increase the distance between each other with age for most regions considered in the study. These results suggest that the median FWVF *f* exhibits a heteroskedastic nature, i.e., the parameter variability increases with age.

In order to evaluate the possible effects from sex and intelligence over the results, we extended the analysis and incorporated (1) the continuous full-scale intelligence quotient variable (*FSIQ*; see Model 2 in Methods) and (2) the binary variable *Sex* (see Model 3 in Methods), and verified the two-way interactions between the variables for both models, i.e. *FSIQ* × *Age* and *Sex* × *Age*. The results show that no main effects from the *FSIQ* and no interactions *FSIQ* × *Age* over all ROIs considered in the study at the significance level of 0.05 except for the CGh with *FSIQ* (*p* < .01) and *FSIQ* × *Age* (*p* < .05) (see Supplementary Table 2). Regarding the gender, we recognize that the variable *Sex* is statistically significant at the level of 0.05 in three regions, namely the ALIC, PCR and CT, all at *p* < .05, and significant interaction term *Sex* × *Age* over two ROIs, i.e. PTR (*p* < .05) and PCR (*p* < .01) (see Supplementary Table 3).

### Free-water volume fraction exhibits heteroskedastic region-specific variations with age

The experiments included in this section illustrate the variability of the FWVF across the adult lifespan and uncover these changes quantitatively. First, in Fig. 2a, we present the region-specific population density plots of the FWVF over the age intervals for the whole WM and seven selected ROIs (see more ROIs in Supplementary Fig. 2). Each density plot aggregates the water fraction values directly from the subjects’ native spaces. Since different regions of the brain are characterized by different sizes and the age intervals are not evenly split in our population sample, the numbers of bins are defined dynamically, being proportional to Doane’s formula (see Methods). The experiment reveals that (1) the population density plots are bell shaped (positively skewed) through all age intervals, (2) the density plots increase the tails and the mode of the density plots shifts positively along successive age decades, and (3) the FWVF values are not affected by the CSF component due to potential image misregistration.

**Figure 2.**
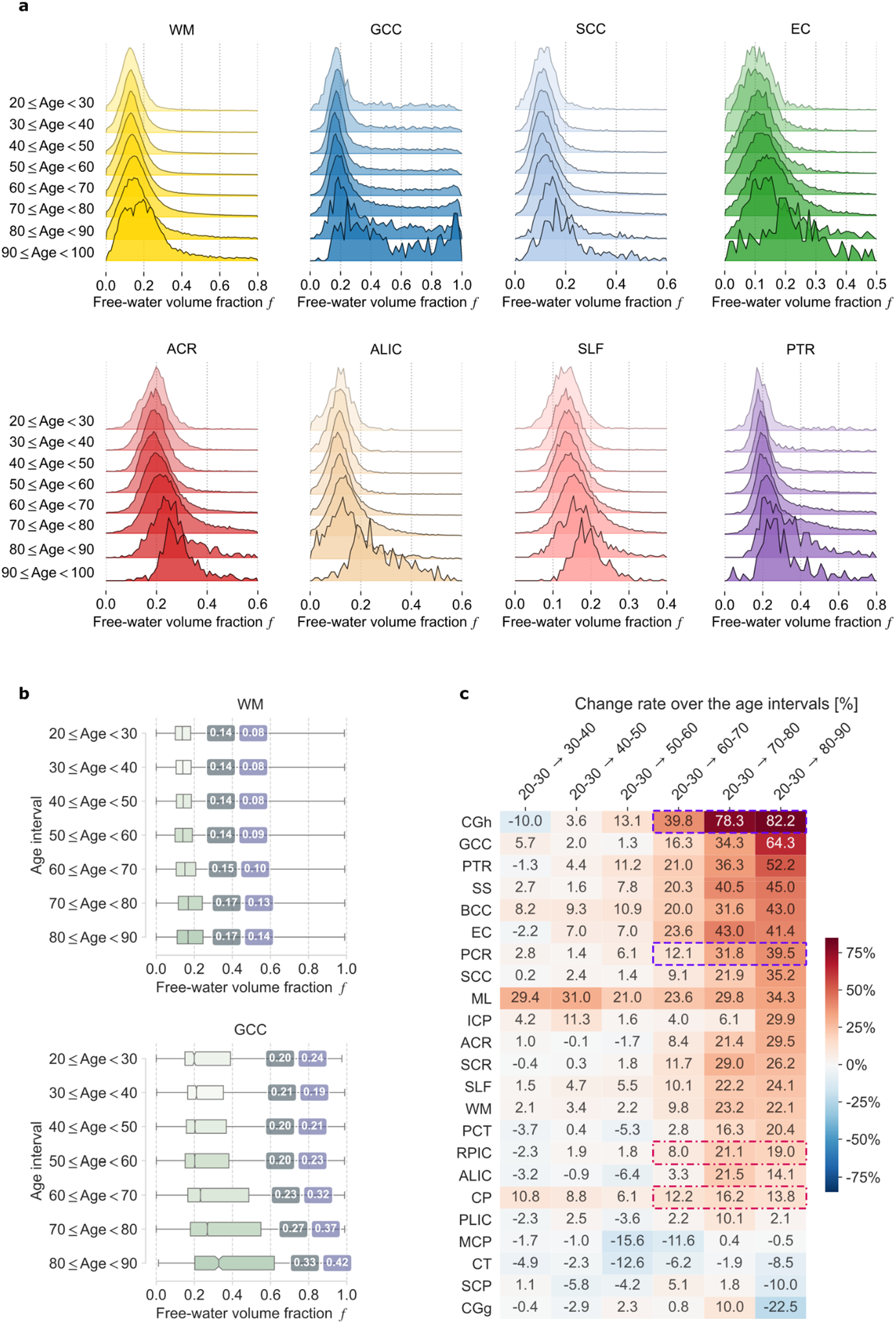
**a** The changes in the population density plots of FWVF *f* presented over the age intervals defined in Table 1 and the eight WM ROIs from Fig. 1b. A single region is represented by eight normalized density plots, each constructed from all subjects’ FWVFs per interval. Doane’s method (Doane, 1976) was applied separately to each ROI and age interval to determine the optimal number of bins used to construct the density plot. The colors of the density plots follow those defined in Fig 1b. **b** The quantitative changes in the FWVF *f* over the WM and GCC and age intervals considered in the study. The box plots present the first, the second (median) and the third quartile, respectively. The whiskers denote minimal and maximal values of the FWVF over the ROIs, while the values over the box plots indicate the population median (first value) and the interquartile range, both calculated from all subjects available per age interval. We have not considered the age interval 90 ≤ *Age* < 100 in this population experiment owing to the presence of only two subjects aged over 90 in the cross-sectional sample. **c** The change rates in the FWVFs over the age intervals are summarized over all ROIs considered in the study. A single number refers to the relative change (in %) of the median FWVF, calculated for a given age interval, with respect to the median FWVF, obtained for the age interval ^2^0 ≤ *Age* < 30. All FWVF’s medians were obtained directly from the subjects’ native spaces. The regions were sorted in descending order according to the values within the interval 80 ≤ *Age* < 90. The regions covered by the same frame colors follow the anterior-posterior spatial gradient (i.e., GCC-SCC and ALIC-PLIC). All population-based experiments report the FWVF variations using only cross-sectional samples.

Next, in Fig. 2b, we report the population FWVF values for the whole WM and the GCC using box plot representation, median values, and interquartile ranges (IRQs) for each age interval (the box plots for remaining ROIs can be found in Supplementary Fig. 3). The experiment reveals that the population median FWVF varies over the WM between 0.14 for base age interval 20 ≤ *Age* < 30 and 0.17 under 80 ≤ *Age* < 90, and it begins to accelerate after the sixth decade of life. Note that this variation increases for other ROIs, like GCC, where it ranges from 0.20 to 0.33. Essentially, the FWVF is a region-specific parameter spanning from 0.09 for the MCP to 0.21 for the SCP region, all reported for the base age interval. Importantly, we observe an increase in the dispersion of the water parameter represented by the IQR measure with age. The regions with a small median initial water fraction (e.g., MCP, ML, EC, CP, SCC, PLIC, ALIC) do not substantially increase their values across the age decades. In contrast, the areas characterized by higher median water fraction values such as the SS, GCC, ACR, PTR or PCR generally show the most substantial parameter growth under the ninth decade of life. In addition, note that these results suggest that the FWVF *f* exhibits a heteroskedastic nature, i.e., the parameter variability increases with age, as indicated by general growth of the IQR in most regions.

**Figure 3.**
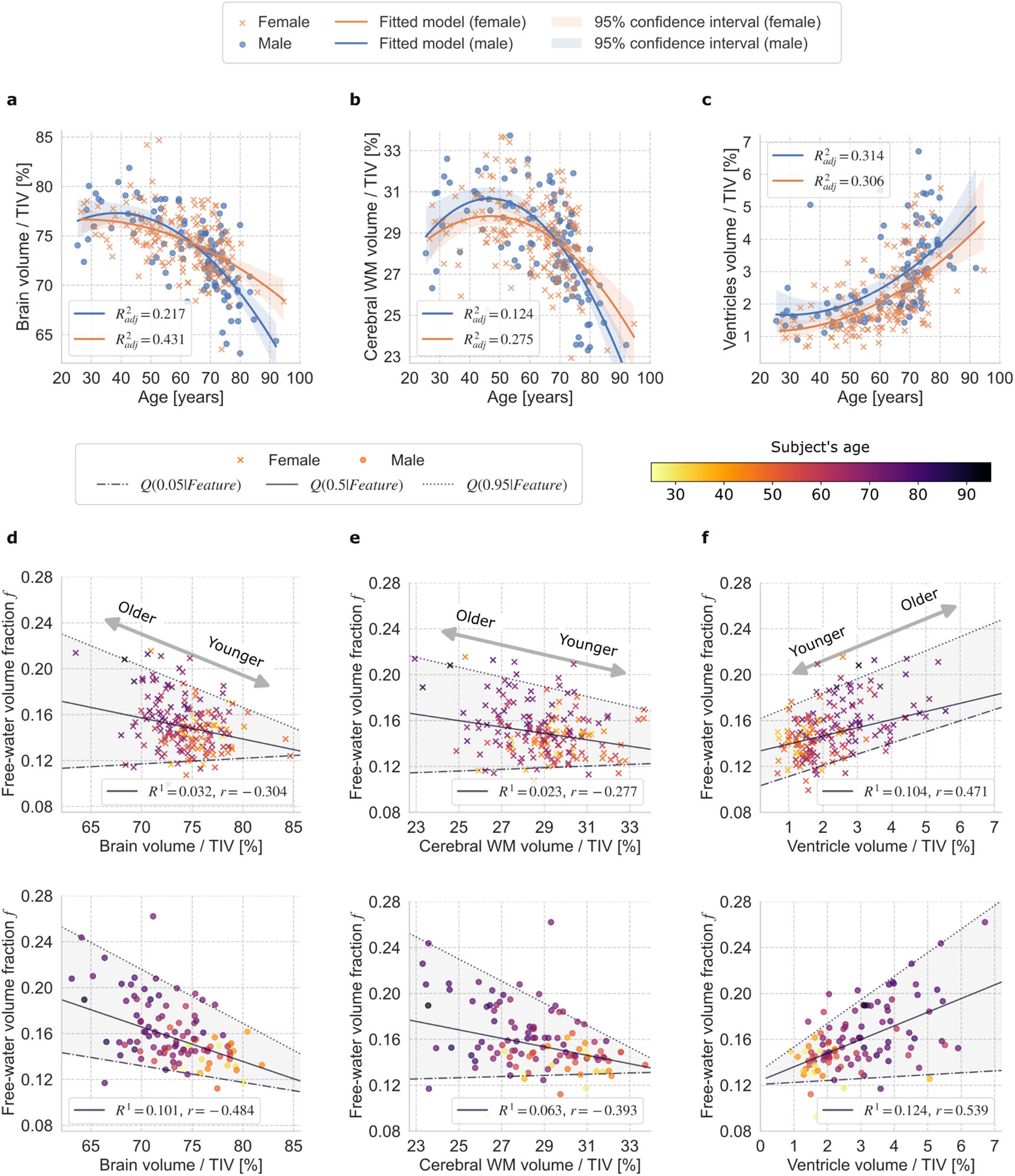
**a-c:** Adult lifespan variations in volumetry-based features of the brain as a function of age (given in years) are presented separately for male and female subjects: **a** brain volume, **b** a cerebral WM volume and **c** ventricles volume. All volume-based features are given in normalized units by the total intracranial volumes (TIV, given in %). The second-order polynomial models were fitted separately for female and male subjects using a linear regression. The adjusted R-squared *R*^2^ parameter characterizes all fitted models. The shaded regions show the 95% confidence intervals estimated from 10,000 bootstrap resamples. **d-e:** The changes in the FWVF *f* as a function of volume-based features (*Feature* ∈ {*Brain volume, Cerebral WM volume, VentriCles volume*}) normalized by the TIV, presented separately for female (top row) and male subjects. An individual marker delivers a median value of the FWVF *f* computed from the WM in the native space of a single subject. The linear QR is used to model the relationship between the FWVF *f* and the volume-based features of the brain: **d** brain volume, **e** cerebral WM volume and **f** ventricles volume. The color of the marker refers to the age of the subject and follows the colorbar included in the right top. The solid lines represent the function *Q*(0.5|*Feature*), the dashed-dotted lines show *Q*(0.05|*Feature*) and the dotted lines represent *Q*(0.95|*Feature*). The regions between *Q*(0.05|*Age*) and *Q*(0.95|*Age*) were shaded for visualization purposes. The goodness-of-fit *R*^1^ at *τ* = 0.5 and Pearson’s correlation coefficient *r* are provided separately for each plot. The arrows show the directions presenting changes in subjects’ age. The experiments employ only cross-sectional samples.

Fig. 2c derives from this experiment and demonstrates the relative errors (given in %) of the median FWVF calculated between the successive intervals and the base interval ^2^0 ≤ *Age* < 30. On average, the WM region is denoted by a relatively small increase in the FWVF until the sixth decade of life, and then it starts enhancing significantly, reaching the value of 22.1% in the ninth decade of life. The highest increase in the water fraction across the life decades has been observed for the CGh (+82.2%) and the GCC (+64.3%). Contrarily, we denote irregular changes in the MCP, CT, SCP and CGg regions with a tendency toward negative values measured at the age interval 80 ≤ *Age* < 90. This experiment also reveals an anterior-posterior spatial gradient of the FWVF, as recently suggested by Chad et al., 2018. More precisely, we observe the higher change rates in the GCC over SCC and ALIC over PLIC (see the color frames included in Fig. 2c).

### Free-water volume fraction shows a non-linear trend with age

In the next experiment, we correlate the FWVF *f* with volumetry-based features, including brain, cerebral WM and ventricles volumes. Prior to that, however, we inspect the changes in the volumetric features for our database and their trajectories as a function of age, to ensure that the behavior matches that previously reported by other authors. Fig. 3a-c presents the variations in these features modeled using the second-order polynomial over the relative values normalized by the total intracranial volume (TIV). The non-normalized values of these features can be seen in Supplementary Fig. 5. In general, we observe the U-shaped characteristics of the whole brain and cerebral WM volumes. The ventricles volume follows a systematic non-linear increase, also accentuating higher values in male subjects. The results obtained with this sanity test are consistent with previous literature reports (Lebel et al., 2012; Bethlehem et al., 2022).

Before studying the correlation between FWVF *f* and the volume-based features, we applied the studentized Breusch-Pagan (BP) test (Breusch and Pagan, 1979; Koenker, 1981) to verify the heteroskedasticity in the FWVF parameter as a function of the anatomical feature. The BP test results were as follows (see Supplementary Table 4):

1. FWVF *f* versus brain volume (females *χ*^2^ (1, *N* = 178) = 12.57, *p* < .001, males *χ*^2^ (1, *N* = 109) = 6.82, *p* = .009);
2. FWVF *f* versus cerebral WM volume (females *χ*^2^ (1, *N* = 178) = 6.41, *p* = .011, males *χ*^2^ (1, *N* = 109) = 6.85, *p* = .009); and
3. FWVF *f* versus ventricles volume (females *χ*^2^ (1, *N* = 178) = 3.78, *p* = .052, males *χ*^2^ (1, *N* = 109) = 24.82, *p* < .001).

These results suggest a heteroskedastic nature of the FWVF parameter as a function of the above-mentioned volume-based features in most cases under the significance level of 0.05. Therefore, the employment of the QR over a standard regression is well founded.

In Fig. 3d-e, we thus relate the FWVF medians calculated over the WM with three volumetry-based features and complement them with the first-order QR models. First, we notice that the linear coefficients for brain and cerebral WM volumes at *τ* = 0.5 are statistically significant under the significance level of 0.05 (see Supplementary Table 4). In contrast, the linear coefficients for ventricles approach the borderline of significance (females *p* = .054, males *p* = .05). For most statistically significant coefficients, we recognize the standard error as at least one order of magnitude smaller than the corresponding coefficient. The results show negative correlations of the FWVF parameter with brain volume (females *r* = −0.304, males *r* = −0.484) and cerebral WM volume (females *r* = −0.277, males *r* = −0.393), while there is a positive correlation between the FWVF and the ventricles volume (females *r* = 0.471, males *r* = 0.539), all under the significance of 0.001. Besides, the experiment reveals stronger correlations for males than for females in all three cases. Apart from the previously confirmed more notable variability of the FWVF with age (see Fig. 1c), here we identify more variable volumes with age that are coupled with higher FWVF values.

### Correcting the DTI representation for the free-water volume fraction leads to flattening trajectories of mean diffusivity and fractional anisotropy measures across the adult lifespan

In this final section, we study the effect of the FWVF on the DTI metrics across the adult lifespan. To that end, we explore the lifespan trajectories of MD and FA measures estimated from the two-component representation given by Eq. (3) and relate them to the standard DTI equivalents, where no FW is assumed. The variations in MD and FA are modeled again using the QR approach presenting now the conditional median of a DTI parameter given the explanatory variable *Age* (see Model 1). For the sake of illustration, in Fig. 4a we visually demonstrate the variations of MD and FA parameters for a randomly chosen male subject at the age of 69. We observe considerable decreased (increased) values of the MD (FA) parameters in the FW compensated DTI scenario. The contours of the WM area become detectable with the FW corrected MD, while the FW compensation generally strengthens the FA measure. In Fig. 4b, we show the population density plots of MD and FA measures for WM, GCC and SCC, calculated from standard and FW compensated DTI representations, depicted over the age intervals previously defined (see more ROIs in Supplementary Fig. 6). The standard MD and FA parameters are typically more dispersed regardless of the age interval, with a positive (negative) shift of the peaks observed for the MD (FA) measure in elderlies. Notice that the population density plots in FW compensated DTI remain nearly unchanged in shape and displacement.

**Figure 4.**
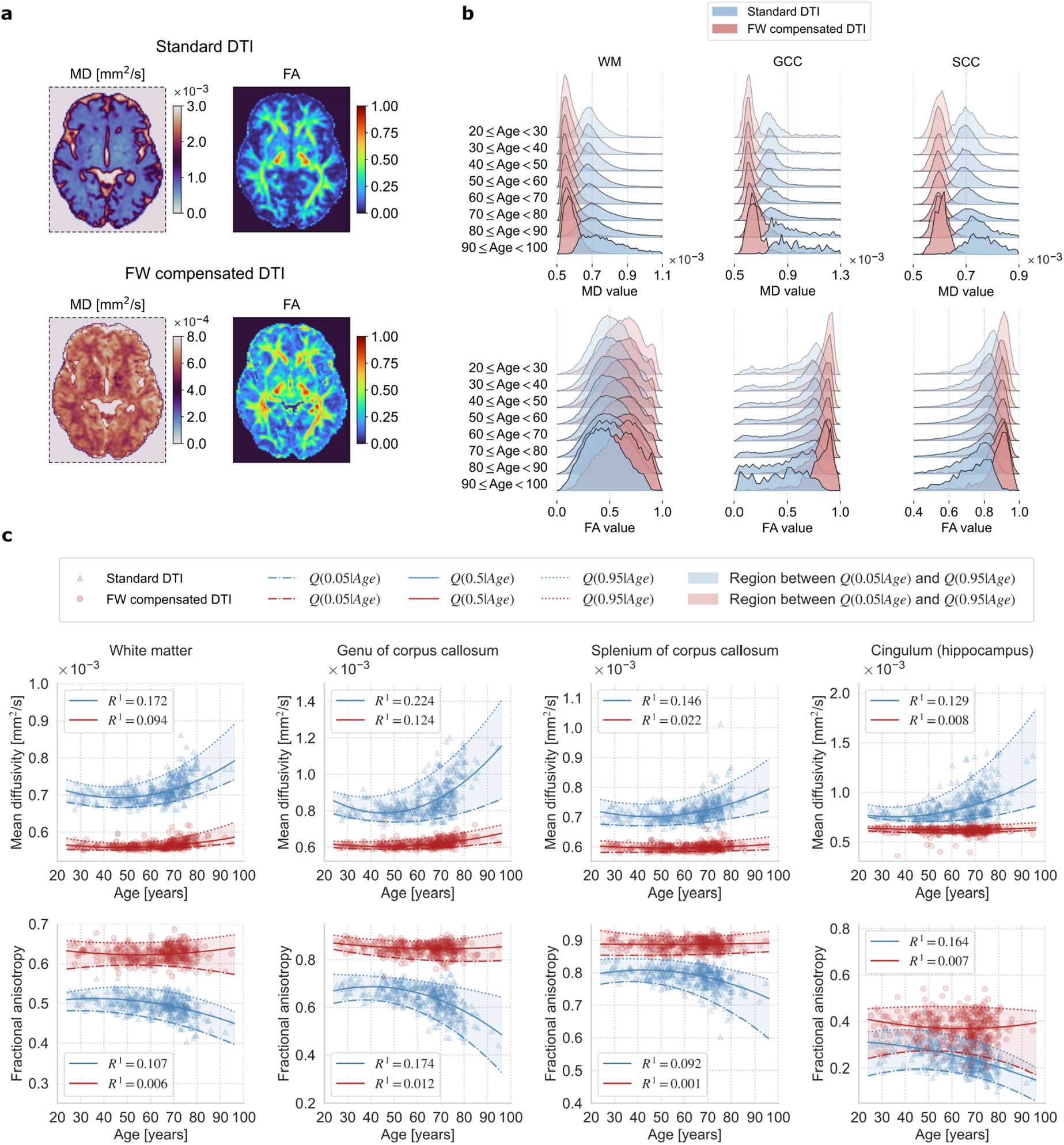
**a** Visual inspection of DTI-based measures based on a standard single-component DTI representation with no FWVF assumption (i.e., *f* = 0) and with a FW compensated DTI. The MD/FA measures were calculated for a randomly chosen healthy male subject at the age of 69 and are shown in a selected axial plane. **b** The density plots presenting the sample variations in MD and FA estimated under the standard DTI and FW compensated DTI over the age intervals defined in Table 1. The experiment reports the qualitative MD and FA variations using only cross-sectional samples. The number of bins for each density plot were determined separately for each age interval using the formula built upon Doane’s method. The population-based experiment uses only cross-sectional samples. **c** Estimated MD/FA measures using a standard DTI and with a FW compensation, and their trajectories across the adult lifespan modeled via the QR technique. The experiment uses cross-sectional and longitudinal samples. Each marker represents the median value of the measure calculated over the ROI in the subject’s native space. The solid lines show the quantile function *Q*(0.5|*Age*), the lower dashed-dotted lines indicate *Q*(0.05|*Age*), and the upper dotted lines present *Q*(0.95|*Age*), all three were computed for a standard DTI (blue lines) and FW compensated DTI (red lines). The regions between *Q*(0.05|*Age*) and *Q*(0.95|*Age*) were shaded for visualization purposes. The goodness-of-fit *R*^1^ at *τ* = 0.5 was computed separately for a standard DTI and FW compensated DTI measures using the procedure introduced by Koenker and Machado (1999).

**Figure 5.**
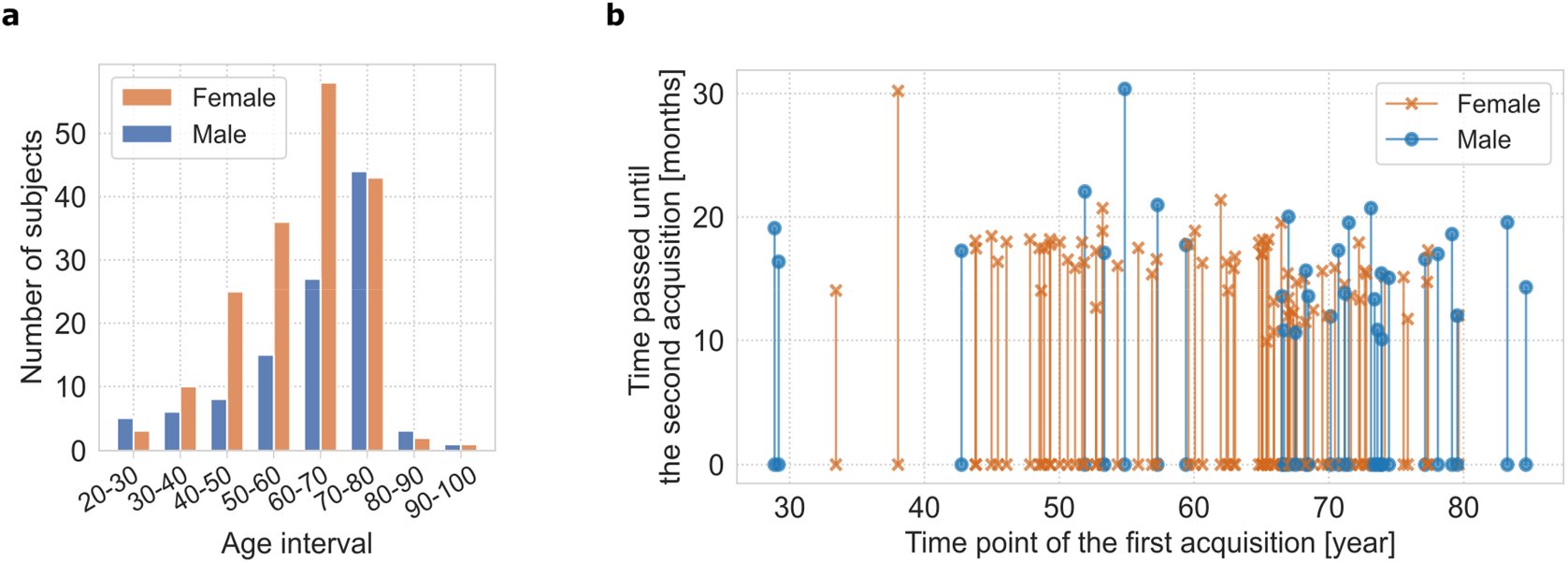
**a** The demography of the study (*N* = 287 healthy cases, 178F/109M) ordered into the age intervals. A single group includes all subjects at age greater or equal to the left range and lower to the right range. **b** Graph presenting the first acquisition time point (in years; horizontal axis) and the time passed until the second acquisition (in months; vertical axis). The second acquisition is available for 99 out of 287 subjects.

Next, in Fig. 4c we represent the changes of MD and FA indices as a function of age for the whole WM and three selected regions, all illustrated with and without the FW compensation (see more ROIs in Supplementary Fig. 7). The coefficients of the models fitted under both variants can be found in Supplementary Table 5 and 6. In general, we observe a non-linear behavior of the standard DTI-based MD parameter over the WM (*p* < .001 for all model coefficients, *R*^1^ = 0.17^2^) with an increasing trend starting from the middle adulthood, and opposite to that for the FA parameter (*p* < .01 for the quadratic model coefficient, *R*^1^ = 0.107), both consistent with previous reports (Westlye et al., 2010; Lebel et al., 2012; Beck et al., 2021). Once the FW correction is applied to the standard DTI, we recognize smaller (larger) values of FW compensated MD (FA) compared to standard DTI, i.e., *p* < .001 for all model coefficients and *R*^1^ = 0.094 for the MD, and non-significant linear (*p* = .196) and quadratic (*p* = .191) model coefficients and *R*^1^ = 0.006 for the FA measure. Besides, the heteroskedastic pattern of the FW corrected DTI measures with age is significantly reduced.

## Discussion

The free-water fraction has become a relevant biomarker in neuroimaging, especially for modeling various types of brain damage (Ofori et al., 2015; Bergamino et al., 2021; Carreira Figueiredo et al., 2022). With its proven importance in age-related pathologies and aging processes (Chad et al., 2018; Edde et al., 2020, Berger et al. 2022), the evolution of this parameter across the adult lifespan has not been studied in detail. Most works that evaluate the microstructural changes in the brain with age based on diffusion MRI simply use a single component DTI approach, where the FWVF is not taken into account (Burzynska et al., 2010; Bennett et al., 2010; Westlye et al., 2010; Lebel et al., 2012; Cox et al., 2016; Beck et al., 2021). The present study investigates precisely how the FWVF varies across the adult lifespan in the healthy human brain WM, as well as its interference with DTI-based metrics. To that end, we employ the spherical means technique (Tristán-Vega et al., 2022) that robustly estimates the water fraction for complex neural architectures, such as fiber crossings, and a two-component dMRI signal representation to retrieve the FW compensated DTI measures.

Although previous studies have already determined that the FWVF changes with age (Metzler-Baddeley et al., 2012; Cox et al., 2016; Chad et al., 2018; Kubicki et al., 2019), in this study, we uncovered a positive trend with age and revealed a non-linear gain in the parameter after the sixth decade of life over most WM areas (Fig. 1c, Supplementary Fig. 1b,d, Supplementary Fig. 3). Our findings suggest a heteroskedastic nature of the FWVF with age that had not been described previously, the existence of a posterior-anterior gradient and the region-specific nature of the changes of the water fraction across the adult lifespan. Additionally, the FWVF showed a linear correlation with volumetric parameters such as the brain volume, cerebral WM volume and ventricles volume (Fig. 3d-f). Finally, we confirmed that the DTI scalar measures do change with age, following a U-shaped trajectory as previously observed by other authors (Westlye et al., 2010; Lebel et al., 2012; Beck et al., 2021). However, in this work, we have detected that those trajectories were deeply influenced by the FW component. Once the FWVF is removed, scalar measures demonstrate a significantly reduced variability over the adult lifespan compared to those calculated from standard DTI (Fig. 4b,c, Supplementary Fig. 6. Supplementary Fig. 7).

The first contribution, then, is to uncover the non-linear behavior and the heteroskedastic nature of the variation of the free-water component with age. As indicated above, this variation of the FWVF with age was already known and it had been previously studied by other authors. However, those studies differ from ours in two main aspects: (1) the regions considered for the study; and (2) the way in which the behavior is modeled. Regarding the former, some studies are limited to the WM skeleton or to specific regions (Metzler-Baddeley et al., 2012; Chad et al., 2018), while our study covers the whole WM in a region-specific manner. Regarding the modeling, our study employs a polynomial approach with a QR optimization to represent the changes in the FWVF as a function of age. This allows us to consider that the variability of the parameters being investigated across the lifespan may change with age, i.e., they follow a heteroskedastic pattern. To our best knowledge, the heteroskedastic modeling of the dMRI parameters has not been properly handled in the literature before.

Some previous endeavors considered a linear approach with homoskedastic assumptions on the FWVF and/or are limited to a selected period of life (Metzler-Baddeley et al., 2012; Chad et al., 2018). Additionally, even when a non-linear model is used to represent the trajectory of FWVF (Kubicki et al., 2021), it is optimized via the non-linear least squares approach where no heteroskedastic assumptions are possible on the dependent variable *Age*. Despite the fact that this polynomial model is well-accepted in the community to account for the variations in diffusion parameters of the brain (Westlye et al., 2010; Beck et al., 2021; Kiely et al., 2022), some studies suggest more *avant-garde* models such as a Poisson curve to account for different changing rates in childhood and aging life stages (Lebel et al., 2012; Kubicki et al., 2019). Nonetheless, compared to our study, these reports used a quadratic cost function in a non-linear fitting procedure and do not allow for modeling the heteroskedastic nature of the parameters, which implicitly assume that the variability of the parameters being investigated across the lifespan does not change with age. Our study provides evidence to suggest that this is not the case. The polynomial approach enables us to handle this phenomenon and to model different quantiles, providing a measure of uncertainty of the parameters. Moreover, the linear QR, contrary to the non-linear models, enabled us to evaluate the significances of the corresponding age-related terms. The current study goes much further and enables us to handle both cross-sectional and longitudinal data. The preceding studies in dMRI parameters alterations with age merely juxtaposed young and older subjects (Burzynska et al., 2010; Bennett et al., 2012), evaluated the cross-sectional data (Westlye et al., 2010; Lebel et al., 2012; Metzler-Baddeley et al., 2012; Cox et al., 2016; Chad et al., 2018; Kiely et al., 2022) or longitudinal measurements (Ofori et al., 2015), with rudimentary studies considering both (Maillard et al., 2019; Beck et al., 2021).

Another major contribution of this work is to unveil how the DTI related metrics are affected by the free water component with age. This part of the study gathers two previously known effects that have been studied separately: (1) the change of diffusion metrics with age: when single-component DTI is considered, diffusion measures in the healthy WM of the human brain present a curvilinear U-form across the lifespan. FA shows a rapid growth in childhood and adolescence and then a systematic decline, while MD reveals the opposite trend (Westlye et al., 2010; Lebel et al., 2012; Beck et al., 2021); (2) the effect of the FWVF on DTI-based measures has also been studied previously (Pasternak et al., 2009; Metzler-Baddeley et al., 2012; Chad et al., 2018; Rydhög et al., 2017; Kubicki et al., 2019), with some preliminary research on differences with age (Metzler-Baddeley et al., 2012; Chad et al., 2018; Kubicki et al., 2021), and under the impact of neurodegenerative diseases (Bergamino et al., 2021; Carreira Figueiredo et al., 2022).

In this work, in order to study the two effects jointly, we examined the variability in the MD and FA across the adult lifespan under two different scenarios: standard DTI and the FW compensated DTI. Our results show that the FW component significantly affects the DTI measures with age. When standard DTI is considered, FA and MD shows the U-form behavior across the lifespan, compatible with findings in Westlye et al. (2010), Lebel et al. (2012), and Beck et al. (2021). On the other hand, when the two-component representation given by Eq. (3) is assumed and the influence of the FW is removed, we observe a flattening effect of the MD and FA measures compared to the standard DTI (see Fig. 4b,c, Supplementary Fig. 6, Supplementary Fig. 7). The flattening effect of the MD parameter has previously been noticed in the fornix with adult subjects (Metzler-Baddeley et al., 2012) and suggested in MD and FA measures over the WM skeleton through a significant decrease in the absolute correlation coefficient with age (Chad et al., 2018). This study has not confirmed the FW corrected FA behavior over the brain WM what others had found in the fornix area (Metzler-Baddeley et al., 2012), i.e., an increase in the elevation of the parameter with age. Instead, we observe a reduced dynamics of the FA measure under a FW compensation (compatible with Chad et al. (2018)) under all regions illustrated except for the SCR, PCR and SCP, reflected in an increase in *R*^1^ parameter. Interestingly, the opposite behavior of the FA measure is observed after the correction, i.e., the non-convex trajectory changes to a convex one (see WM and cingulum (hippocampus) in Fig. 4c). The variability reduction in DTI-based measures under the FW compensation has been previously observed by Albi et al. (2017) and explained as an increase in reproducibility. However, our results explain that the FW correction reduces the variability of the MD/FA measures across lifespan but does not necessarily imply an improvement in their reproducibility.

Different authors have studied the variation of the FA across the lifespan without considering the FWVF and, therefore, they have tried to explain this effect differently. Vernooij et al. (2008) explained the age-related decrease in FA by atrophy and lesion formation. This hypothesis was rejected by Westlye et al. 2010, where authors pointed out that such pathological factors cannot have a significant impact on healthy young subjects. Alternatively, they hypothesized that the formation of redundant myelin and water compartments may increase volume and decrease FA with age. Bennett et al. (2010), Burzynska et al. (2010) and Lebel et al. (2012) also relate FA alterations with changes of myelination and/or axonal density. Our experiments have shed some light over the interpretation of the results: the origins of these variations may be strongly associated with the FWVF, although other biological effects must also be taken into account.

All in all, our results indicate that the changes in the FW component constitute a major factor to explain the variability of DTI metrics with age. This issue must be taken into account when using these metrics for clinical studies. Depending on the ages of the participants, data can present some bias due to the lifespan related changes. Fortunately, if the dMRI acquisition comprises at least two different shells, the estimation of the FWVF is feasible and it is possible to correct the age-related bias. In addition, when comparing scalar measures like FA and MD in healthy and pathological subjects, it is tempting to explain the differences between groups as the result of underlying biological effects: the reduction of myelin, changes of the axonal density and so on. In the light of our experiments, we should be more cautious in explaining microstructural processes from the results obtained from diffusion measures, since there may be unconsidered factors at play, such as the FWVF.

This study presents several limitations that must be noted. First, a relatively small sample size was employed (*N* = 287), and the distribution of the samples across the age intervals is non-uniform. However, compared to the previous studies modeling the free-water fraction variations across the lifespan (Metzler-Baddeley et al., 2012; Cox et al., 2016; Chad et al., 2018), our population sample has been spread over the whole adult life, including subjects from early adulthood until senescence. Moreover, the sample includes a mixed cross-sectional and longitudinal design that enables us to model the FWVF over the lifespan, including changes over time, suppressing the effect of inter-subject variability. Secondly, as with most image-based population studies, the current study is limited by the registration procedure. The approach used in our study follows the well-accepted standard community pipeline, i.e., we non-linearly register the FA to the FSL template “FMRIB58_FA” and then inversely warp the atlas ROIs to the subjects’ native spaces using the FSL software suite. A proper registration procedure and template selection are critical for ROI-based studies as the misalignment might bias the representative values. Recently, (Wu et al., 2022) proposed a new FA template that meets the requirements of lifespan studies and reduces possible misalignment artifacts in elderly brains. Here, however, each ROI had been morphologically eroded before all analyses took place to exclude potential outliers exposed to the partial volume effect. In fact, the ROI-based analysis might be considered a deficiency as such. Nonetheless, some previous lifespan studies in diffusion MRI (Tamnes et al., 2010; Kiely et al., 2022) follow this line since it is standardized, as mentioned above.

Additionally, we limited the scope of the water component to the simplest form of diffusion representation, DTI, although one can extrapolate the study and incorporate more advanced dMRI signal representations such as diffusion kurtosis imaging (DKI) (Jensen et al., 2005) or the propagator-based metrics (Özarslan et al., 2013; Tristán-Vega and Aja-Fernández, 2021). For instance, the DKI includes information beyond the standard DTI appearing at a higher b-value regime (Jensen et al., 2005), while the propagator-based measures can catch the axial or planar diffusion profiles (Özarslan et al., 2013). The proper water-correction of these dMRI signal representations might lead us to new insights into modeling the non-Gaussian diffusion profiles and unbiased indicators of restrictive diffusion and cellularity across the lifespan.

Finally, another limitation of the study is related to the method made use of to estimate the FWVF. The method in Tristán-Vega et al. (2022) adapts the general formulation from Tristán-Vega and Aja-Fernández (2021) by fixing the parallel diffusivity in Eq. (2) over the white matter area to *λ*_par_ = 2.1 · 10^−3^ mm^2^/s. This way, the FWVF can be estimated from just two different acquired shells. However, as recently shown by Tristán-Vega and Aja-Fernández (2021), the parameter *λ*_par_ can be constant over the WM tissue without the repercussions of further inferences. Guerrero et al. (2019) has additionally shown that the parallel diffusivity used in the NODDI does not vary appreciably with age unless the population group includes infants or adolescent subjects and has no sex effects. Finally, we notice, as the FW and restricted water follow different signal decays, that they can be separated using dedicated MRI machines equipped with high-strength gradients (Afzali et al., 2022). However, using ultra-strong gradients (up to 300 mT/m) exceeds the current technical capabilities in versatile population studies, imposing a clinical scanner with a common gradient strength, as about 40 mT/m employed in this study.

Modern and successful brain aging research requires more than standard group-based comparisons, and new directions related to trajectory-based tracking of diseases are expected to gain interest in the near future. This raises new challenges in the neuroimaging field with a particular emphasis on capturing the onset of neurodegenerative disease and predicting further alterations of the brain. This paper lays the foundations for the analysis of the evolution of FWVF in the healthy human brain WM across the lifespan changes, also gaining insight on the effects of the compensation of DTI measures for FW in these analyses.

## Acknowledgements

Tomasz Pieciak, Guillem París, Antonio Tristán-Vega, Rodrigo de Luis-García and Santiago Aja-Fernández acknowledge Ministerio de Ciencia e Innovación Gobierno de España with research grants PID2021-124407NB-I00. Tomasz Pieciak acknowledges the Polish National Agency for Academic Exchange for grant PPN/BEK/2019/1/00421 under the Bekker programme and the Ministry of Science and Higher Education (Poland) under the scholarship for outstanding young scientists (692/STYP/13/2018). Guillem París was funded by the Consejería de Educación de Castilla y León and the European Social Fund through the “Ayudas para financiar la contratación predoctoral de personal investigador - Orden EDU/1100/2017 12/12” program. The study is supported by the Research Council of Norway (223273, 249795, 298646, 300767), the South-Eastern Norway Regional Health Authority (2014097, 2019101), the Norwegian ExtraFoundation for Health and Rehabilitation (2015/FO5146), KG Jebsen Stiftelsen, and the European Research Council under the European Union’s Horizon 2020 research and Innovation program (ERC 802998).

