## Supplementary materials for "Free-water volume fraction increases non-linearly with age in the white matter of the healthy human brain"

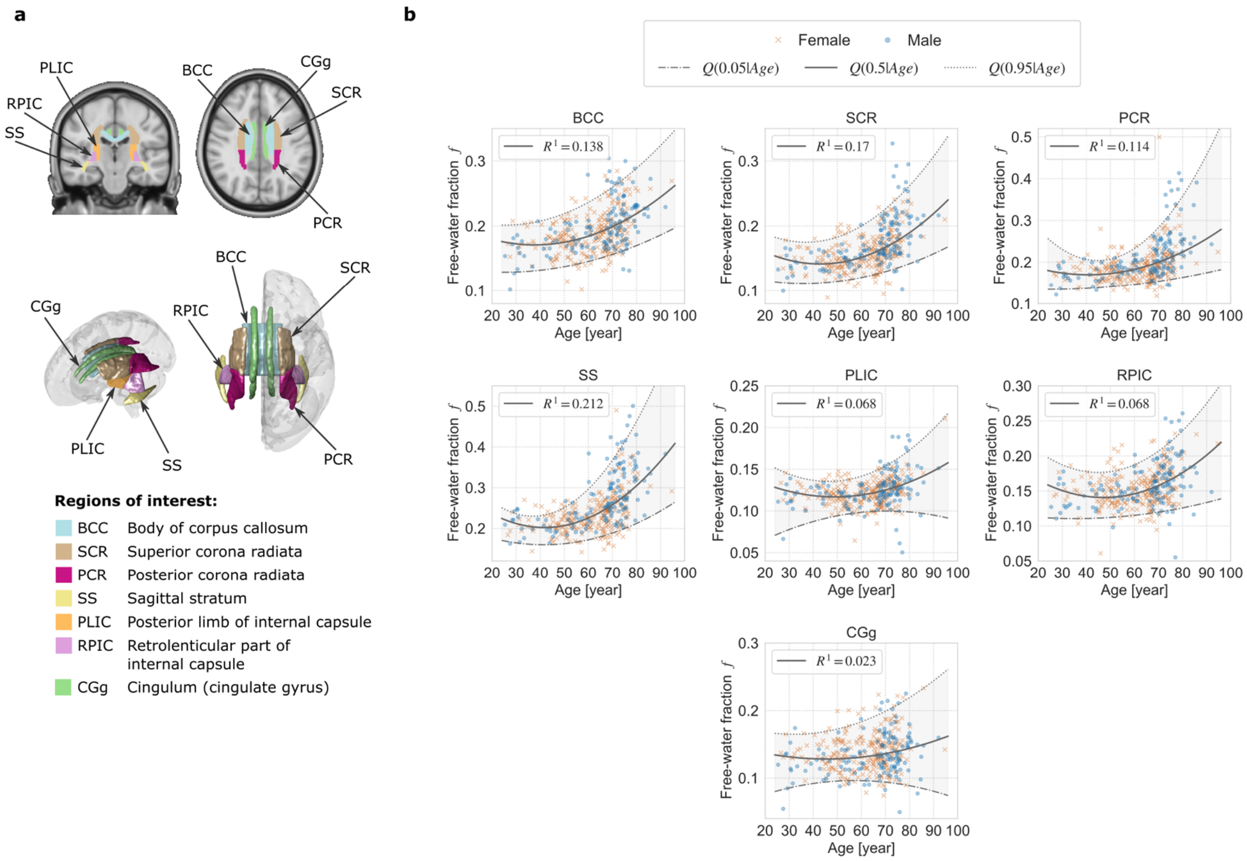


**Supplementary Figure 1.** The changes of the FWVF across the lifespan over different ROIs. **a,c** The regions of interest (ROIs) used in the study were retrieved from the JHU DTI-based atlas (Mori et al., 2005) and visualized in the standard space over the T1-weighted coronal and axial slices, and using the 3D model. **b,d** Estimated FWVFs $f$ and their trajectories computed with a quantile regression technique via a polynomial fitting over different ROIs presented in panels **a** and **c**. Each marker delivers a median value of the FWVF calculated for a single subject over the ROI in the native space. The experiment uses cross-sectional and longitudinal samples. The quantile function $Q(\tau|Age)$ for a single ROI is shown under three quantiles, i.e., the solid line presents $Q(\tau|Age)$ under $\tau = 0.5$ (50th percentile), the lower thin dashed-dotted line indicates $Q(\tau|Age)$ under $\tau = 0.05$ (5th percentile), and the upper thin dotted line presents $Q(\tau|Age)$ under $\tau = 0.95$ (95th percentile), all three computed from median FWVFs. The regions between $Q(0.05|Age)$ and $Q(0.95|Age)$ were shaded for visualization purposes. The goodness-of-fit $R^1$ at $\tau = 0.5$ was computed for each region using the procedure introduced by Koenker and Machado (1999).

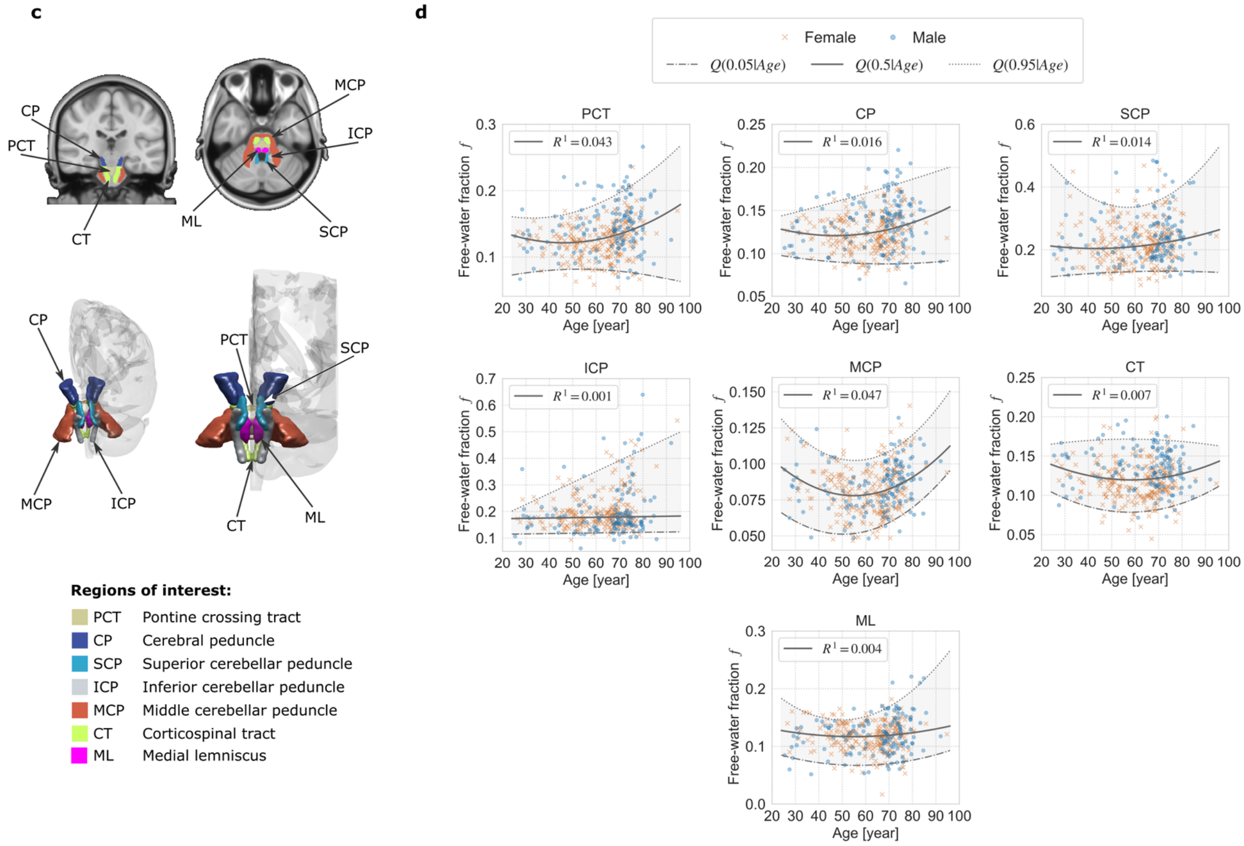


**Supplementary Figure 1. (continued)**

**Supplementary Table 1.** The coefficients of Model 1 (see Methods) fitted to the FWVF median values from Fig. 1c and Supplementary Fig. 1b,d using the QR technique under the quantile $\tau = 0.5$, including all ROIs considered in the study. The columns $\beta_0$, $\beta_1$ and $\beta_2$ represent the constant, linear ($Age$) and quadratic ($Age^2$) terms of the model, respectively. The standard errors of the coefficients are given in the parentheses. The *p*-values refer to the significance of the coefficients: ****p* < .001, ***p* < .01, **p* < .05. If the coefficient is non-significant under the significance level of 0.05 the cell is marked in red and the *p*-value is given in the superscript. The column $R^1(\tau)$ includes the goodness-of-fit of the model at $\tau = 0.5$ (Koenker and Machado, 1999). The next to last column presents the Akaike information criterion (AIC) (Akaike, 1974) of the second-order model (AIC 2nd order) versus the first-order model (AIC 1st order). The smaller the AIC, the better the model represents the FWVF changes across the lifespan. The smaller AIC of the two models are bolded. The last column includes age peaks derived from quadratic models and are given in years (y) and months (m). The row names follow the abbreviations defined in Fig. 1b and Supplementary Fig. 1a,c.

| ROI | Quantile regression fitting coefficients, *p*-values, standard errors (std. err.), $R^1(\tau)$ at $\tau = 0.5$, AIC criterions and age peaks. | | | | | |
|---|---|---|---|---|---|---|
| | $\beta_0$ $^{p\text{-value}}$ (std. err.) | $\beta_1$ $^{p\text{-value}}$ $\times 10^2$ (std. err. $\times 10^2$) | $\beta_2$ $^{p\text{-value}}$ $\times 10^3$ (std. err. $\times 10^3$) | $R^1(\tau)$ | AIC 2nd order (AIC 1st order) | Age peak |
| WM | 0.206333*** (0.018275) | -0.288098*** (0.058608) | 0.030973*** (0.004711) | 0.141 | **-1874.99** (-1834.48) | 46y 6m |
| GCC | 0.403430*** (0.050253) | -0.935097*** (0.191771) | 0.102469*** (0.015918) | 0.192 | **-1174.39** (-1111.16) | 45y 8m |
| SCC | 0.182367*** (0.019278) | -0.314751*** (0.072667) | 0.034894*** (0.005083) | 0.134 | **-1731.82** (-1692.77) | 45y 1m |
| CGh | 0.249897*** (0.075467) | -0.586569 $^{p = .053}$ (0.290605) | 0.080598** (0.026210) | 0.14 | **-760.43** (-744.77) | 36y 5m |
| EC | 0.155080*** (0.020770) | -0.248037** (0.091385) | 0.032443*** (0.008036) | 0.132 | **-1543.68** (-1524.76) | 38y 3m |
| ACR | 0.295054*** (0.025157) | -0.457579*** (0.099203) | 0.050790*** (0.008427) | 0.142 | **-1419.5** (-1382.34) | 45y 1m |
| ALIC | 0.232953*** (0.024894) | -0.479542*** (0.109031) | 0.047800*** (0.009790) | 0.085 | **-1589.05** (-1554.23) | 50y 2m |
| SLF | 0.192256*** (0.019674) | -0.244965** (0.082501) | 0.027468*** (0.006670) | 0.09 | **-1666.78** (-1647.91) | 44y 7m |
| PTR | 0.199782*** (0.032785) | -0.104364 $^{p = .398}$ (0.152529) | 0.025355 $^{p = .079}$ (0.014096) | 0.157 | **-1236.69** (-1229.3) | 20y 7m |

**Supplementary Table 1. (continued)**

| ROI | Quantile regression fitting coefficients, *p*-values, standard errors (std. err.), $R^1(\tau)$ at $\tau = 0.5$, AIC criterions and age peaks. | | | | | |
|---|---|---|---|---|---|---|
| | $\beta_0$ $^{p\text{-value}}$ (std. err.) | $\beta_1$ $^{p\text{-value}}$ $\times 10^2$ (std. err. $\times 10^2$) | $\beta_2$ $^{p\text{-value}}$ $\times 10^3$ (std. err. $\times 10^3$) | $R^1(\tau)$ | AIC 2nd order (AIC 1st order) | Age peak |
| BCC | 0.209508*** (0.021841) | -0.203675* (0.076463) | 0.026961*** (0.007006) | 0.138 | **-1596.31** (-1579.73) | 37y 9m |
| SCR | 0.205971*** (0.027413) | -0.300001** (0.087521) | 0.034973*** (0.007844) | 0.17 | **-1700.82** (-1664.93) | 42y 11m |
| PCR | 0.229869*** (0.030470) | -0.292918* (0.114158) | 0.035802** (0.010641) | 0.114 | **-1441.09** (-1417.33) | 40y 11m |
| SS | 0.324931*** (0.030757) | -0.587173*** (0.120127) | 0.070249*** (0.011908) | 0.212 | **-1282.81** (-1231.8) | 41y 10m |
| PLIC | 0.161161*** (0.016496) | -0.180773* (0.068223) | 0.018468*** (0.005314) | 0.068 | **-2081.63** (-2060.16) | 48y 11m |
| RPIC | 0.214897*** (0.030141) | -0.314075** (0.102602) | 0.033203*** (0.008344) | 0.068 | **-1695.5** (-1675.93) | 47y 4m |
| CGg | 0.155654*** (0.014729) | -0.120458* (0.059509) | 0.013241* (0.005073) | 0.023 | **-1619.23** (-1615.02) | 45y 6m |
| PCT | 0.170942*** (0.021314) | -0.214398** (0.082999) | 0.023215** (0.007831) | 0.043 | **-1556.94** (-1544.4) | 46y 2m |
| CP | 0.151702*** (0.028151) | -0.130857 $^{p=.140}$ (0.080118) | 0.013901* (0.007852) | 0.016 | **-1711.42** (-1707.69) | 47y 1m |
| SCP | 0.243593*** (0.064660) | -0.183922 $^{p=.336}$ (0.215646) | 0.021360 $^{p=.210}$ (0.019784) | 0.014 | **-921.74** (-919.34) | 43y 1m |
| ICP | 0.170346*** (0.014106) | 0.013025 $^{p=.551}$ (0.020012) | - | 0.001 | -994.32 (**-995.98**) | - |
| MCP | 0.140166*** (0.022786) | -0.226118*** (0.070870) | 0.020541*** (0.005767) | 0.047 | **-2019.09** (-1985.83) | 55y 1m |
| CT | 0.176855*** (0.034500) | -0.196321 $^{p=.064}$ (0.121762) | 0.016833 $^{p=.076}$ (0.009970) | 0.007 | **-1689.88** (-1686.73) | 58y 4m |
| ML | 0.150156*** (0.030301) | -0.120727 $^{p=.249}$ (0.118803) | 0.010956 $^{p=.247}$ (0.010356) | 0.004 | **-1619.57** (-1619.37) | 55y 1m |

**Supplementary Table 2.** The coefficients of Model 2 (see Methods) fitted to the FWVF median values from Fig. 1c and Supplementary Fig. 1b,d using the QR technique under the quantile $\tau = 0.5$, including all ROIs considered in the study. The columns $\beta_0$, $\beta_1$ and $\beta_2$ represent the constant, linear ($Age$) and quadratic ($Age^2$) terms of the model, while $\beta_3$ and $\beta_4$ refer to the variable $FSIQ$ and the interaction between $FSIQ$ and $Age$, respectively. The standard errors of the coefficients are given in the parentheses. The *p*-values refer to the significance of the coefficients: ****p* < .001, ***p* < .01, **p* < .05. If the coefficient is non-significant under the significance level of 0.05 the cell is marked in red and the *p*-value is given in the coefficient superscript. The column $R^1$ includes the goodness-of-fit of the model at $\tau = 0.5$ (Koenker and Machado, 1999). The next to last column presents the Akaike information criterion (AIC) (Akaike, 1974) of the second-order model (AIC 2nd order) versus the first-order model (AIC 1st order). The smaller the AIC, the better the model represents the FWVF changes across the lifespan. The smaller AIC of the two models are bolded. The row names follow the abbreviations defined in Fig. 1b and Supplementary Fig. 1a,c.

| ROI | Quantile regression fitting coefficients, *p*-values, standard errors (std. err.), $R^1(\tau)$ at $\tau = 0.5$ and AIC criterions. | | | | | | |
|---|---|---|---|---|---|---|---|
| | $\beta_0^{p\text{-value}}$ (std. err.) | $\beta_1^{p\text{-value}} \times 10^2$ (std. err. $\times 10^2$) | $\beta_2^{p\text{-value}} \times 10^3$ (std. err. $\times 10^3$) | $\beta_3^{p\text{-value}} \times 10^2$ (std. err. $\times 10^2$) | $\beta_4^{p\text{-value}} \times 10^4$ (std. err. $\times 10^4$) | $R^1$ | AIC 2nd order (AIC 1st order) |
| WM | 0.227384** (0.083147) | -0.275877* (0.143490) | 0.029772*** (0.005829) | $-0.021030^{p=.765}$ (0.069322) | $0.001299^{p=.991}$ (0.122574) | 0.146 | **-1875.16** (-1841.64) |
| GCC | $0.300652^{p=.095}$ (0.170202) | -0.668290* (0.336796) | 0.109027*** (0.013183) | $0.099766^{p=.462}$ (0.155608) | $-0.282106^{p=.222}$ (0.253153) | 0.206 | **-1183.2** (-1108.49) |
| SCC | 0.233550** (0.076950) | -0.363220** (0.114052) | 0.034736*** (0.006923) | $-0.044479^{p=.543}$ (0.077176) | $0.044365^{p=.708}$ (0.123235) | 0.137 | **-1730.09** (-1693.78) |
| CGh | $-0.286890^{p=.123}$ (0.192822) | $0.238578^{p=.578}$ (0.397253) | 0.091851*** (0.028363) | 0.493225** (0.160129) | -0.817365* (0.343310) | 0.149 | **-765.32** (-744.04) |
| EC | $0.075047^{p=.408}$ (0.094943) | $-0.043122^{p=.805}$ (0.162892) | 0.031067** (0.009678) | $0.064327^{p=.384}$ (0.073220) | $-0.160477^{p=.259}$ (0.130703) | 0.143 | **-1549.86** (-1528.54) |
| ACR | 0.373651** (0.126002) | -0.539117* (0.216617) | 0.053501*** (0.010074) | $-0.062976^{p=.500}$ (0.103962) | $0.047920^{p=.763}$ (0.180494) | 0.146 | **-1419.06** (-1381.43) |
| ALIC | $0.201609^{p=.052}$ (0.097776) | -0.400476* (0.199237) | 0.048888*** (0.010852) | $0.028720^{p=.750}$ (0.089701) | $-0.076756^{p=.608}$ (0.132440) | 0.091 | **-1590.0** (-1554.99) |
| SLF | 0.267575*** (0.082200) | -0.311437* (0.144721) | 0.024879*** (0.006996) | $-0.076746^{p=.287}$ (0.063540) | $0.090663^{p=.455}$ (0.114901) | 0.095 | **-1667.02** (-1651.75) |
| PTR | $0.153790^{p=.194}$ (0.117134) | $0.064444^{p=.788}$ (0.264511) | $0.023875^{p=.070}$ (0.014541) | $0.033288^{p=.743}$ (0.104958) | $-0.126093^{p=.538}$ (0.200766) | 0.162 | **-1237.24** (-1231.5) |

**Supplementary Table 2. (continued)**

| ROI | Quantile regression fitting coefficients, ***p***-values, standard errors (std. err.), $R^1(\tau)$ at $\tau = 0.5$ and AIC criterions. | | | | | | |
|---|---|---|---|---|---|---|---|
| | $\beta_0$ $^{p\text{-value}}$ (std. err.) | $\beta_1$ $^{p\text{-value}}$ $\times 10^2$ (std. err. $\times 10^2$) | $\beta_2$ $^{p\text{-value}}$ $\times 10^3$ (std. err. $\times 10^3$) | $\beta_3$ $^{p\text{-value}}$ $\times 10^2$ (std. err. $\times 10^2$) | $\beta_4$ $^{p\text{-value}}$ $\times 10^4$ (std. err. $\times 10^4$) | $R^1$ | AIC 2nd order (AIC 1st order) |
| BCC | 0.211564* (0.086463) | -0.171351 $^{p=.290}$ (0.134598) | 0.027345** (0.007919) | -0.002966 $^{p=.969}$ (0.074625) | -0.027757 $^{p=.817}$ (0.115946) | 0.143 | **-1596.94** (-1583.65) |
| SCR | 0.198662** (0.081003) | -0.237819 $^{p=.059}$ (0.149363) | 0.035531*** (0.008339) | 0.005045 $^{p=.929}$ (0.064900) | -0.053805 $^{p=.587}$ (0.101643) | 0.18 | **-1706.57** (-1673.14) |
| PCR | 0.272175*** (0.079656) | -0.283460 $^{p=.093}$ (0.145759) | 0.035851** (0.013208) | -0.039713 $^{p=.649}$ (0.091688) | -0.002822 $^{p=.985}$ (0.160208) | 0.121 | **-1443.08** (-1422.64) |
| SS | 0.443654*** (0.109036) | -0.777581*** (0.204916) | 0.069960*** (0.010305) | -0.103903 $^{p=.229}$ (0.092961) | 0.167659 $^{p=.299}$ (0.155726) | 0.214 | **-1280.5** (-1230.56) |
| PLIC | 0.186689** (0.060943) | -0.197760 $^{p=.055}$ (0.109740) | 0.019698** (0.005607) | -0.019900 $^{p=.696}$ (0.055406) | 0.004686 $^{p=.952}$ (0.075935) | 0.074 | **-2082.26** (-2062.13) |
| RPIC | 0.267767** (0.077763) | -0.322744* (0.164056) | 0.024563** (0.008715) | -0.072095 $^{p=.344}$ (0.081661) | 0.099105 $^{p=.488}$ (0.127783) | 0.071 | **-1693.53** (-1677.75) |
| CGg | 0.123346 $^{p=.129}$ (0.080951) | -0.048529 $^{p=.744}$ (0.164442) | 0.014883* (0.006212) | 0.031132 $^{p=.640}$ (0.070894) | -0.075693 $^{p=.575}$ (0.130851) | 0.024 | **-1616.05** (-1612.4) |
| PCT | 0.171085* (0.077040) | -0.231911 $^{p=.115}$ (0.176066) | 0.022425** (0.007645) | -0.000282 $^{p=.997}$ (0.074092) | 0.019790 $^{p=.854}$ (0.117380) | 0.044 | **-1553.28** (-1542.28) |
| CP | 0.191123* (0.097938) | -0.172981 $^{p=.291}$ (0.159251) | 0.013471 $^{p=.153}$ (0.009736) | -0.035534 $^{p=.630}$ (0.077264) | 0.041183 $^{p=.744}$ (0.127684) | 0.018 | **-1708.61** (-1706.09) |
| SCP | 0.346421 $^{p=.098}$ (0.211977) | -0.384558 $^{p=.230}$ (0.326498) | 0.014927 $^{p=.497}$ (0.018931) | -0.100549 $^{p=.629}$ (0.213715) | 0.225464 $^{p=.502}$ (0.325725) | 0.018 | **-920.76** (-920.46) |
| ICP | 0.174209 $^{p=.199}$ (0.133746) | 0.091127 $^{p=.703}$ (0.213302) | - | -0.005815 $^{p=.964}$ (0.132246) | -0.064172 $^{p=.752}$ (0.198642) | 0.005 | -993.81 (**-994.84**) |
| MCP | 0.067211 $^{p=.432}$ (0.084911) | -0.150343 $^{p=.208}$ (0.119070) | 0.020035*** (0.005840) | 0.062859 $^{p=.448}$ (0.080890) | -0.062945 $^{p=.634}$ (0.115104) | 0.051 | **-2018.65** (-1991.82) |
| CT | 0.193065* (0.101744) | -0.230144 $^{p=.178}$ (0.162788) | 0.021963* (0.009811) | -0.000679 $^{p=.994}$ (0.086996) | -0.020536 $^{p=.870}$ (0.132511) | 0.009 | **-1687.88** (-1683.81) |
| ML | 0.108160 $^{p=.270}$ (0.098149) | -0.001177 $^{p=.994}$ (0.163279) | - | 0.006379 $^{p=.939}$ (0.072970) | 0.005973 $^{p=.960}$ (0.125692) | 0.003 | -1616.21 (**-1616.85**) |

**Supplementary Table 3.** The coefficients of Model 3 (see Methods) fitted to the FWVF median values from Fig. 1c and Supplementary Fig. 1b,d using the QR technique under the quantile $\tau = 0.5$, including all regions of interest considered in the study. The columns $\beta_0$, $\beta_1$ and $\beta_2$ represent the constant, linear ($Age$) and quadratic ($Age^2$) terms of the model, while $\beta_3$ and $\beta_4$ refer to the variable $Sex$ and the interaction between $Sex$ and $Age$, respectively. The standard errors of the coefficients are given in the parentheses. The *p*-values refer to the significance of the coefficients: ****p* < .001, ***p* < .01, **p* < .05. If the coefficient is non-significant under the significance level of 0.05 the cell is marked in red and the *p*-value is given in the coefficient superscript. The column $R^1$ includes the goodness-of-fit of the model at $\tau = 0.5$ (Koenker and Machado, 1999). The next to last column presents the Akaike information criterion (AIC) (Akaike, 1974) of the second-order model (AIC 2nd order) versus the first-order model (AIC 1st order). The smaller the AIC, the better the model represents the FWVF changes across the lifespan. The smaller AIC of the two models are bolded. The row names follow the abbreviations defined in Fig. 1b and Supplementary Fig. 1a,c.

| ROI | Quantile regression fitting coefficients, *p*-values, standard errors (std. err.), $R^1(\tau)$ at $\tau = 0.5$ and AIC criterions. | | | | | | |
|---|---|---|---|---|---|---|---|
| | $\beta_0$ $^{p\text{-value}}$ (std. err.) | $\beta_1$ $^{p\text{-value}}$ $\times 10^2$ (std. err. $\times 10^2$) | $\beta_2$ $^{p\text{-value}}$ $\times 10^3$ (std. err. $\times 10^3$) | $\beta_3$ $^{p\text{-value}}$ $\times 10^1$ (std. err. $\times 10^1$) | $\beta_4$ $^{p\text{-value}}$ $\times 10^3$ (std. err. $\times 10^3$) | $R^1$ | AIC 2nd order (AIC 1st order) |
| WM | 0.189698*** (0.020598) | -0.258458*** (0.065771) | 0.030494*** (0.005008) | 0.186928 $^{p=.145}$ (0.135442) | -0.313983 $^{p=.099}$ (0.215050) | 0.146 | **-1874.9** (-1835.8) |
| GCC | 0.361282*** (0.054870) | -0.852168*** (0.189612) | 0.099965*** (0.015416) | 0.421876 $^{p=.166}$ (0.314039) | -0.698605 $^{p=.167}$ (0.527922) | 0.195 | **-1172.64** (-1113.13) |
| SCC | 0.163552*** (0.026581) | -0.238511*** (0.081209) | 0.029115*** (0.006201) | 0.069988 $^{p=.666}$ (0.159331) | -0.234842 $^{p=.359}$ (0.242564) | 0.143 | **-1735.45** (-1707.13) |
| CGh | 0.209750* (0.088252) | -0.434855 $^{p=.199}$ (0.343757) | 0.069518* (0.031445) | 0.250742 $^{p=.573}$ (0.448199) | -0.655528 $^{p=.449}$ (0.799303) | 0.142 | **-758.48** (-748.49) |
| EC | 0.156860*** (0.027709) | -0.271883* (0.104836) | 0.034742*** (0.009604) | 0.109830 $^{p=.448}$ (0.167559) | -0.060310 $^{p=.809}$ (0.290187) | 0.139 | **-1546.69** (-1523.38) |
| ACR | 0.279259*** (0.035642) | -0.435064*** (0.122387) | 0.051018*** (0.008768) | 0.202397 $^{p=.318}$ (0.200576) | -0.310614 $^{p=.319}$ (0.357051) | 0.144 | **-1417.07** (-1380.73) |
| ALIC | 0.196827*** (0.033481) | -0.394912*** (0.122952) | 0.043372*** (0.009759) | 0.329184* (0.149535) | -0.499055 $^{p=.066}$ (0.249690) | 0.094 | **-1592.63** (-1553.64) |
| SLF | 0.195466*** (0.027317) | -0.259664** (0.094378) | 0.028709*** (0.007469) | -0.000363 $^{p=.998}$ (0.180721) | 0.022087 $^{p=.934}$ (0.260292) | 0.09 | **-1663.24** (-1644.8) |
| PTR | 0.156385*** (0.035197) | 0.049507 $^{p=.729}$ (0.130724) | 0.015328 $^{p=.199}$ (0.012771) | 0.329671 $^{p=.137}$ (0.229059) | -0.856170* (0.384960) | 0.173 | **-1247.44** (-1246.07) |

**Supplementary Table 3. (continued)**

| ROI | Quantile regression fitting coefficients, *p*-values, standard errors (std. err.), $R^1(\tau)$ at $\tau = 0.5$ and AIC criterions. | | | | | | |
|---|---|---|---|---|---|---|---|
| | $\beta_0$ $^{p\text{-value}}$ (std. err.) | $\beta_1$ $^{p\text{-value}} \times 10^2$ (std. err. $\times 10^2$) | $\beta_2$ $^{p\text{-value}} \times 10^3$ (std. err. $\times 10^3$) | $\beta_3$ $^{p\text{-value}} \times 10^1$ (std. err. $\times 10^1$) | $\beta_4$ $^{p\text{-value}} \times 10^3$ (std. err. $\times 10^3$) | $R^1(\tau)$ | AIC 2nd order (AIC 1st order) |
| BCC | 0.188166*** (0.027249) | -0.145562 $^{p=.130}$ (0.094978) | 0.023334** (0.009116) | 0.167882 $^{p=.259}$ (0.161268) | -0.291078 $^{p=.201}$ (0.259929) | 0.142 | **-1596.26** (-1582.85) |
| SCR | 0.181499*** (0.023037) | -0.241863** (0.076709) | 0.032917*** (0.006832) | 0.245329 $^{p=.107}$ (0.136137) | -0.475489 $^{p=.052}$ (0.235402) | 0.179 | **-1704.93** (-1667.82) |
| PCR | 0.206340*** (0.032978) | -0.242565* (0.120868) | 0.035037** (0.011714) | 0.427972* (0.210483) | -0.835737** (0.337436) | 0.134 | **-1454.2** (-1437.74) |
| SS | 0.286914*** (0.046436) | -0.464854*** (0.142851) | 0.062301*** (0.012974) | 0.280351 $^{p=.195}$ (0.215876) | -0.619444 $^{p=.095}$ (0.390735) | 0.225 | **-1291.72** (-1251.9) |
| PLIC | 0.156686*** (0.019116) | -0.172843* (0.068310) | 0.018135*** (0.005322) | 0.037532 $^{p=.727}$ (0.101540) | -0.045690 $^{p=.781}$ (0.159002) | 0.069 | **-2078.55** (-2057.16) |
| RPIC | 0.214381*** (0.041304) | -0.311792* (0.125978) | 0.032957*** (0.010082) | 0.017514 $^{p=.935}$ (0.208982) | -0.014132 $^{p=.967}$ (0.357969) | 0.069 | **-1691.62** (-1679.47) |
| CGg | 0.134425*** (0.023079) | -0.090302 $^{p=.191}$ (0.075246) | 0.012812* (0.006598) | 0.202649 $^{p=.116}$ (0.138908) | -0.218864 $^{p=.341}$ (0.233103) | 0.034 | **-1624.22** (-1619.1) |
| PCT | 0.161284*** (0.023730) | -0.144918 $^{p=.102}$ (0.082751) | 0.017597* (0.007728) | 0.014689 $^{p=.926}$ (0.170302) | -0.255403 $^{p=.262}$ (0.262426) | 0.072 | **-1576.64** (-1570.09) |
| CP | 0.143640*** (0.026336) | -0.073901 $^{p=.425}$ (0.088978) | 0.009676 $^{p=.139}$ (0.007043) | 0.015839 $^{p=.917}$ (0.140334) | -0.245529 $^{p=.288}$ (0.224492) | 0.054 | **-1738.01** (-1737.34) |
| SCP | 0.237679*** (0.065031) | -0.163668 $^{p=.447}$ (0.244556) | 0.020751 $^{p=.267}$ (0.016967) | 0.211062 $^{p=.583}$ (0.409226) | -0.426749 $^{p=.456}$ (0.629798) | 0.016 | **-919.3** (-917.62) |
| ICP | 0.246444** (0.070883) | -0.300250 $^{p=.238}$ (0.240665) | 0.026632 $^{p=.329}$ (0.025345) | -0.156779 $^{p=.637}$ (0.345125) | 0.548396 $^{p=.335}$ (0.537069) | 0.014 | **-1000.1** (-998.79) |
| MCP | 0.140054*** (0.019012) | -0.243047*** (0.077153) | 0.023083*** (0.006580) | 0.233733 $^{p=.066}$ (0.121266) | -0.369201 $^{p=.056}$ (0.197317) | 0.056 | **-2022.11** (-1992.28) |
| CT | 0.173110*** (0.018597) | -0.101269 $^{p=.152}$ (0.076095) | 0.006605 $^{p=.262}$ (0.006789) | -0.274319* (0.143487) | 0.099233 $^{p=.659}$ (0.225571) | 0.082 | **-1746.99** (-1746.1) |
| ML | 0.143979*** (0.029987) | -0.142279 $^{p=.167}$ (0.101256) | 0.015587 $^{p=.095}$ (0.009097) | 0.282861 $^{p=.160}$ (0.206320) | -0.431611 $^{p=.206}$ (0.341279) | 0.01 | **-1620.64** (-1617.69) |

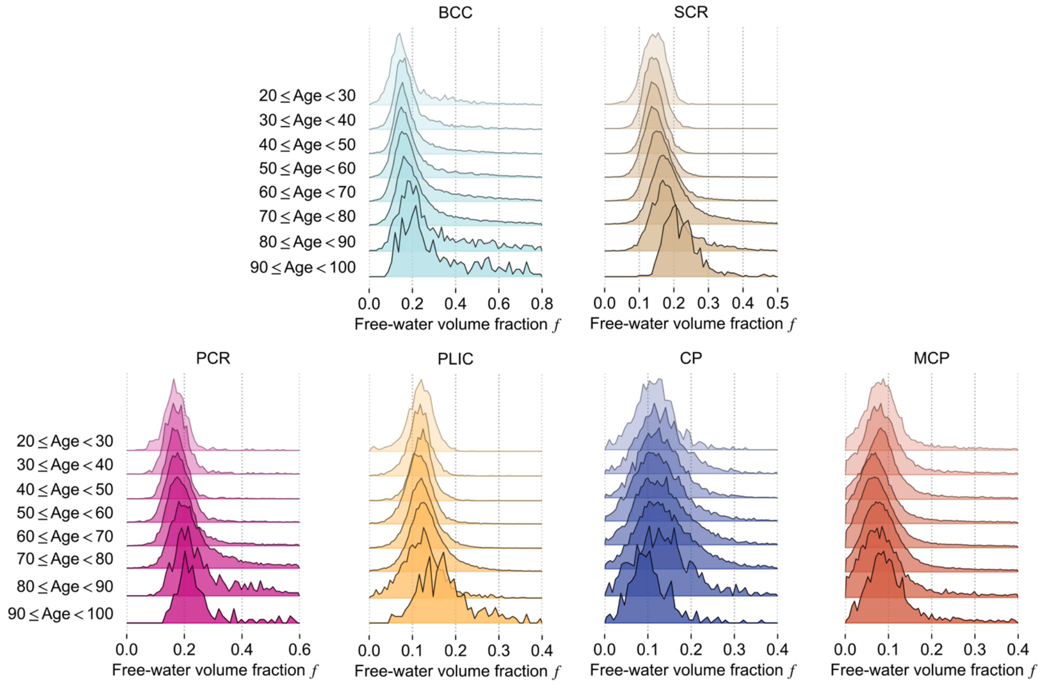


**Supplementary Figure 2.** The changes in the density plots of FWVF $f$ presented across the age intervals defined in Table 1 and brain regions visualized Supplementary Fig. 1a,c. A single region is characterized by eight density plots, each computed from all subjects' water fraction values available per age interval. The FWVFs were aggregated directly from subjects' native spaces to calculate a single density plot over the ROI. The experiment employs only cross-sectional samples. The formula built upon Doane's method (Doane, 1976) was applied separately to each ROI and age interval to determine the optimal number of bins used to construct the density plot (see Methods). All density plots histograms were normalized. For clarity, the colors of all density plots follow those defined in Fig. 1b and Supplementary Fig. 1a,c.

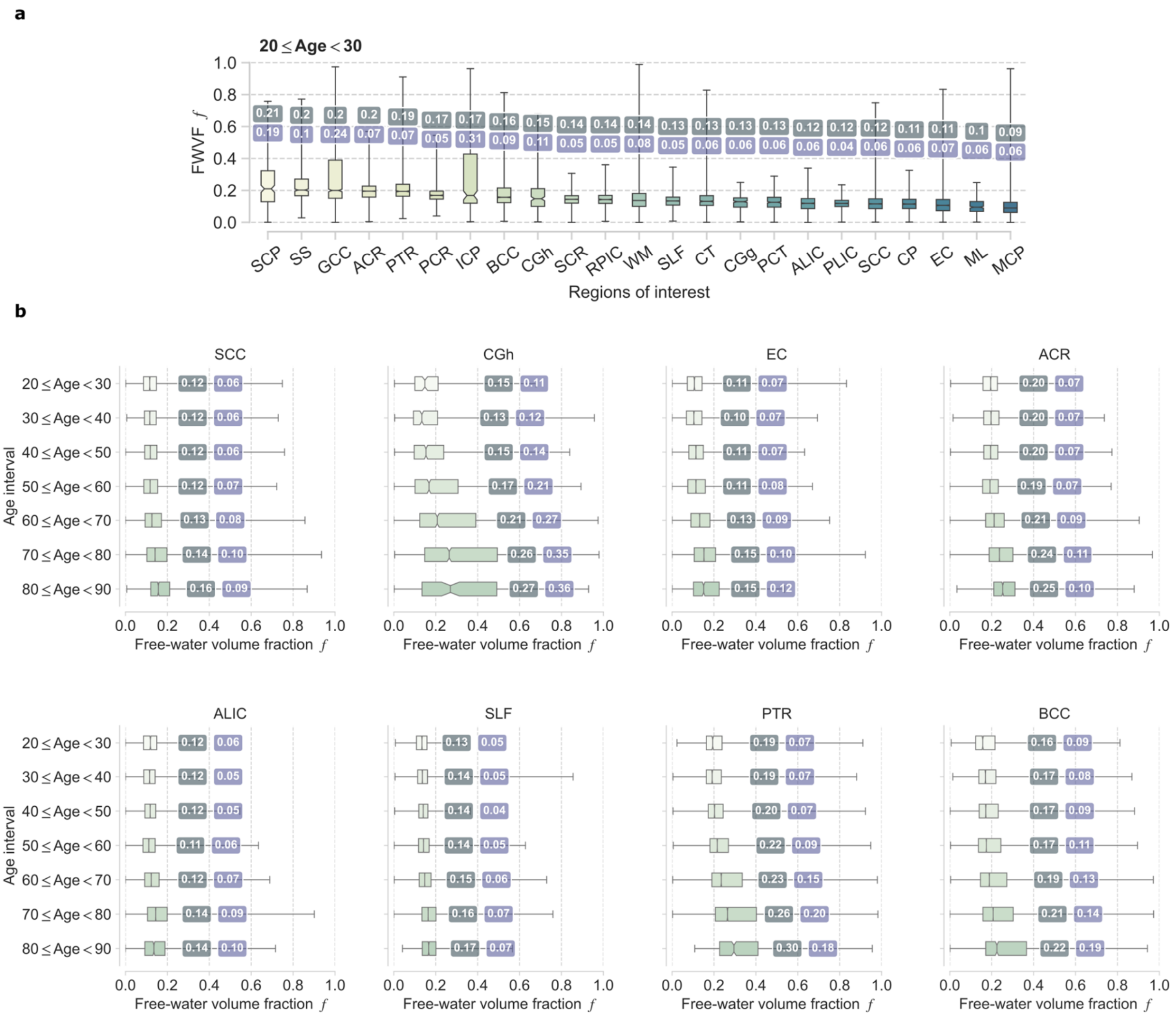


**Supplementary Figure 3.** The quantitative changes in the population FWVF parameter: **a** over the ROIs under the base interval $20 \leq Age < 30$ and **b,c** over the age intervals defined for each ROI separately. The box plots present the first, the second (median) and the third quartile, respectively. The whiskers denote minimal and maximal values of the FWVF, while the values over the box plots indicate the population median (first value) and the interquartile range, calculated from all subjects available per age interval. The experiment reports the quantitative FWVF variations using only cross-sectional samples aggregated from subjects' native spaces. The labels use the abbreviations defined in Fig. 1b and Supplementary Fig. 1a,c. We have not considered the age interval $90 \leq Age < 100$ in this population experiment owing to the presence of only two subjects aged over 90 in the cross-sectional sample.

**c**

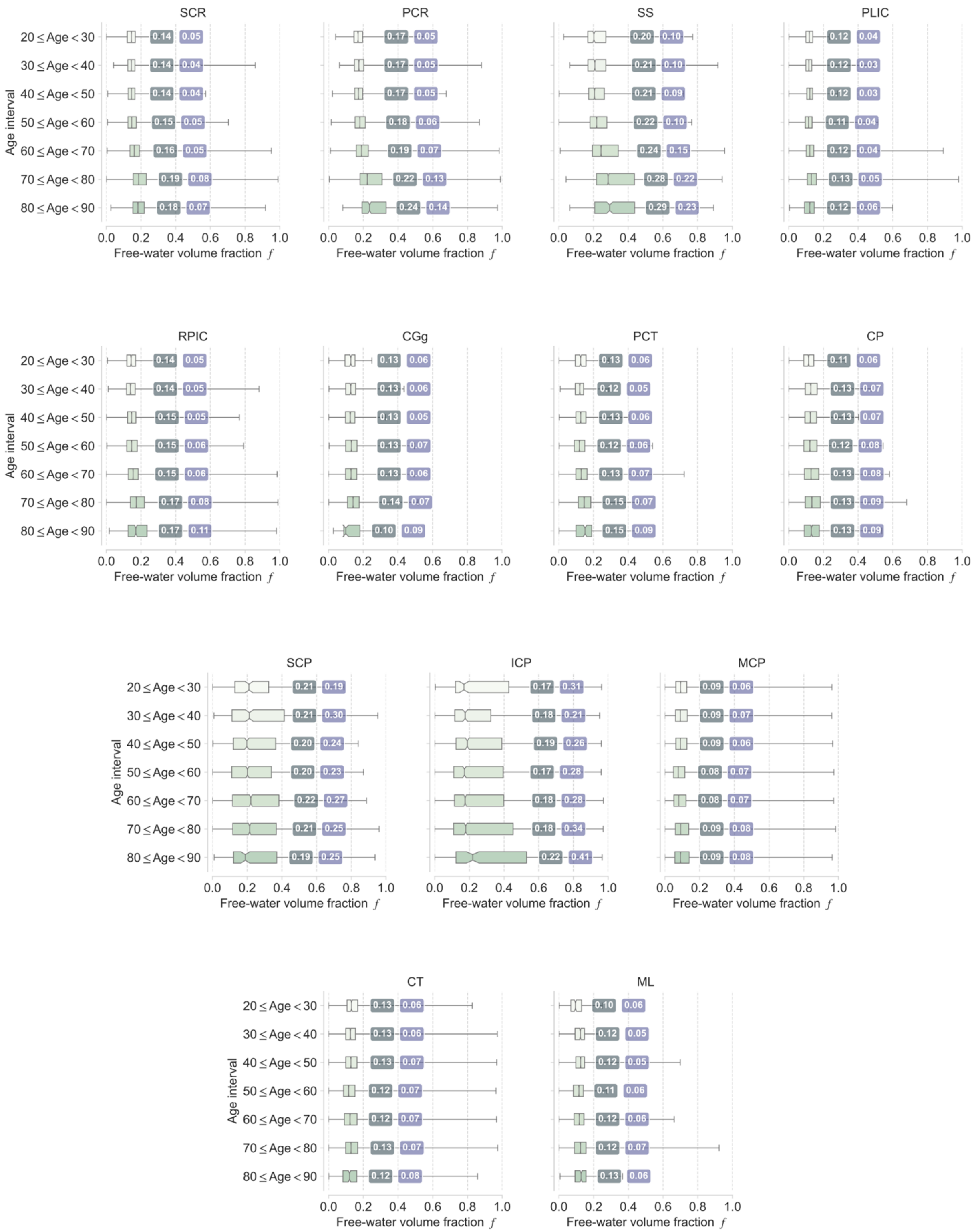


**Supplementary Figure 3. (continued)**

The experiment included in Supplementary Fig. 4 relates the changing rate of the FWVF f (given in %), defined as the relative increase from the third to the ninth decade of life (shown in vertical axes), to the values obtained at the third decade of life (horizontal axes). We observe that the smaller FWVF values are characterized by more prominent and variable changing rates, obeying region-specific differentiae.

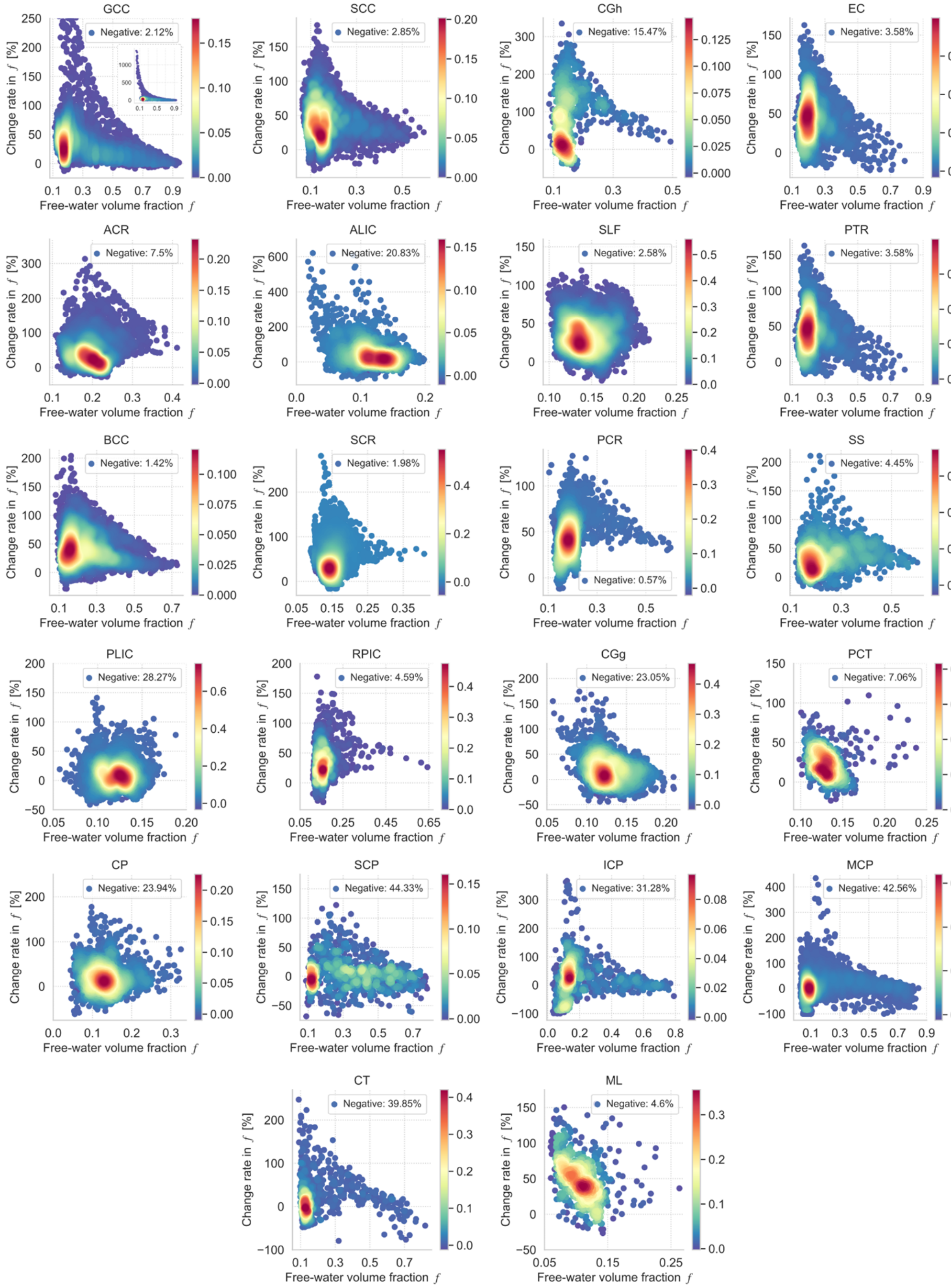


**Supplementary Figure 4.** The 2D density plots illustrating the change rate (i.e. the relative error) of FWVF at the age interval $80 \leq Age < 90$ to the base interval at $20 \leq Age < 30$ as a function of the FWVF at the base interval. The experiment reports the FWVF variations using only cross-sectional samples. All 2D density plots were constructed from the FWVFs warped to the standard space and then interpolated using the bivariate splines. The percentage given for each plot refers to the negative values of the change rate. The labels use the abbreviations defined in Fig. 1b and Supplementary Fig. 1a,c.

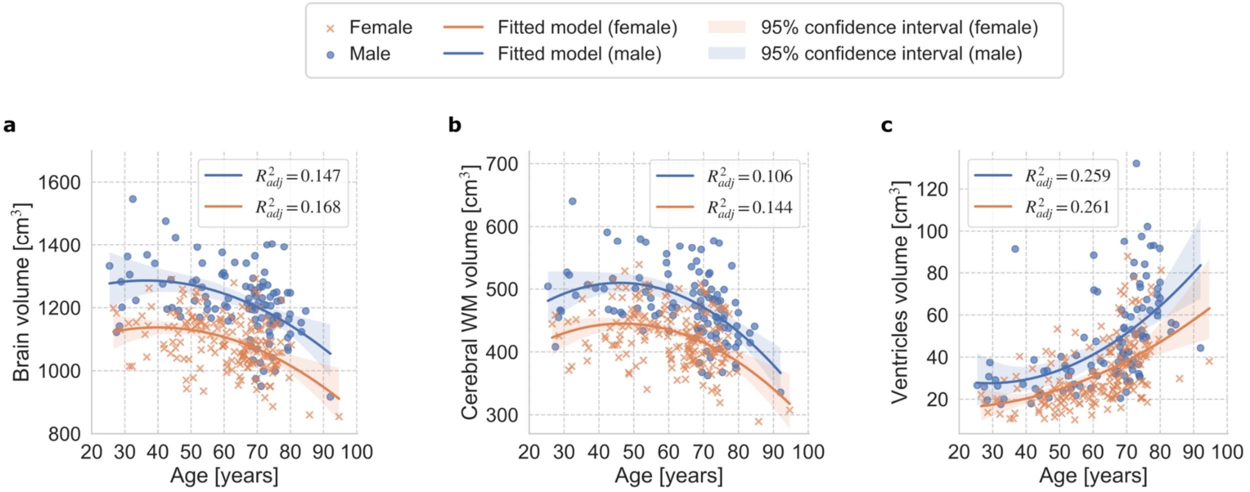


**Supplementary Figure 5.** Adult lifespan variations in volumetry-based features of the brain as a function of age (given in [years]) presented separately for male and female subjects: **a** brain volume, **b** cerebral WM volume and **c** ventricles volume. All volume-based features are absolute values given in [$cm^3$]. The experiment reports the volumetry-based features using only cross-sectional samples. The second-order polynomial models were fitted separately for female (orange lines) and male subjects (blue lines) using a linear regression. The adjusted R-squared $R^2_{adj}$ parameter characterizes all fitted models. The shaded regions show the 95% confidence intervals estimated from 10,000 bootstrap resamples.

**Supplementary Table 4.** The coefficients of Model 4 (see Methods) fitted to the FWVF median values from Fig. 3 using the first-order QR technique under the quantile $\tau = 0.5$, including three volume-based brain features (i.e., brain volume, cerebral WM volume, ventricles volume). Top rows present the fitted models for female subjects, while the bottom refer to the fitted models for male subjects. The columns $\beta_0$ and $\beta_1$ represent the constant and linear ($Feature$) terms of the model, respectively. The standard errors of the coefficients are given in the parentheses and the *p*-values of the coefficients are provided in the superscript. Notation used for the *p*-values: ***$p < .001$, **$p < .01$, *$p < .05$. If the coefficient is non-significant under the significance level of 0.05 the cell is marked in red and the *p*-value is given in the superscript. The column $R^1(\tau)$ includes the goodness-of-fit of the model at $\tau = 0.5$ (Koenker and Machado, 1999), the column Pearson's $r$ presents the correlation coefficient between the FWVF parameter and the feature, and the *p*-value, while the last column refers to the statistics $\chi^2$ and *p*-value obtained from the studentized Breusch-Pagan test (Breusch and Pagan, 1979; Koenker, 1981) used to verify the heteroskedasticity in the FWVF parameter as a function of volume-based feature.

| **Feature** | **Quantile regression fitting coefficients, *p*-values, standard errors (std. err.), $R^1(\tau)$ at $\tau = 0.5$, Pearson's $r$ and *p*-values, the statistics $\chi^2$ and *p*-values from the Breusch-Pagan test.** | | | | |
|---|---|---|---|---|---|
| | $\beta_0^{p\text{-value}}$ (std. err.) | $\beta_1^{p\text{-value}} \times 10^2$ (std. err. $\times 10^2$) | $R^1(\tau)$ | Pearson's $r$ (*p*-value) | Breusch-Pagan test statistics $\chi^2$ (*p*-value) |
| Brain volume | 0.285747*** (0.055513) | -0.183684* (0.074237) | 0.032 | -0.304 ($p < .001$) | 12.57 ($p < .001$) |
| | 0.376143*** (0.061770) | -0.300597*** (0.081792) | 0.101 | -0.484 ($p < .001$) | 6.82 ($p = .009$) |
| Cerebral WM volume | 0.229463*** (0.041466) | -0.278217 $^{p = .054}$ (0.143350) | 0.023 | -0.277 ($p < .001$) | 6.41 ($p = .011$) |
| | 0.260601*** (0.057580) | -0.369270 $^{p = .05}$ (0.186582) | 0.063 | -0.393 ($p < .001$) | 6.85 ($p = .009$) |
| Ventricles volume | 0.132378*** (0.003632) | 0.124106*** (0.008343) | 0.104 | 0.471 ($p < .001$) | 3.78 ($p = .052$) |
| | 0.711147*** (0.160521) | 1.191486** (0.395463) | 0.124 | 0.539 ($p < .001$) | 24.82 ($p < .001$) |

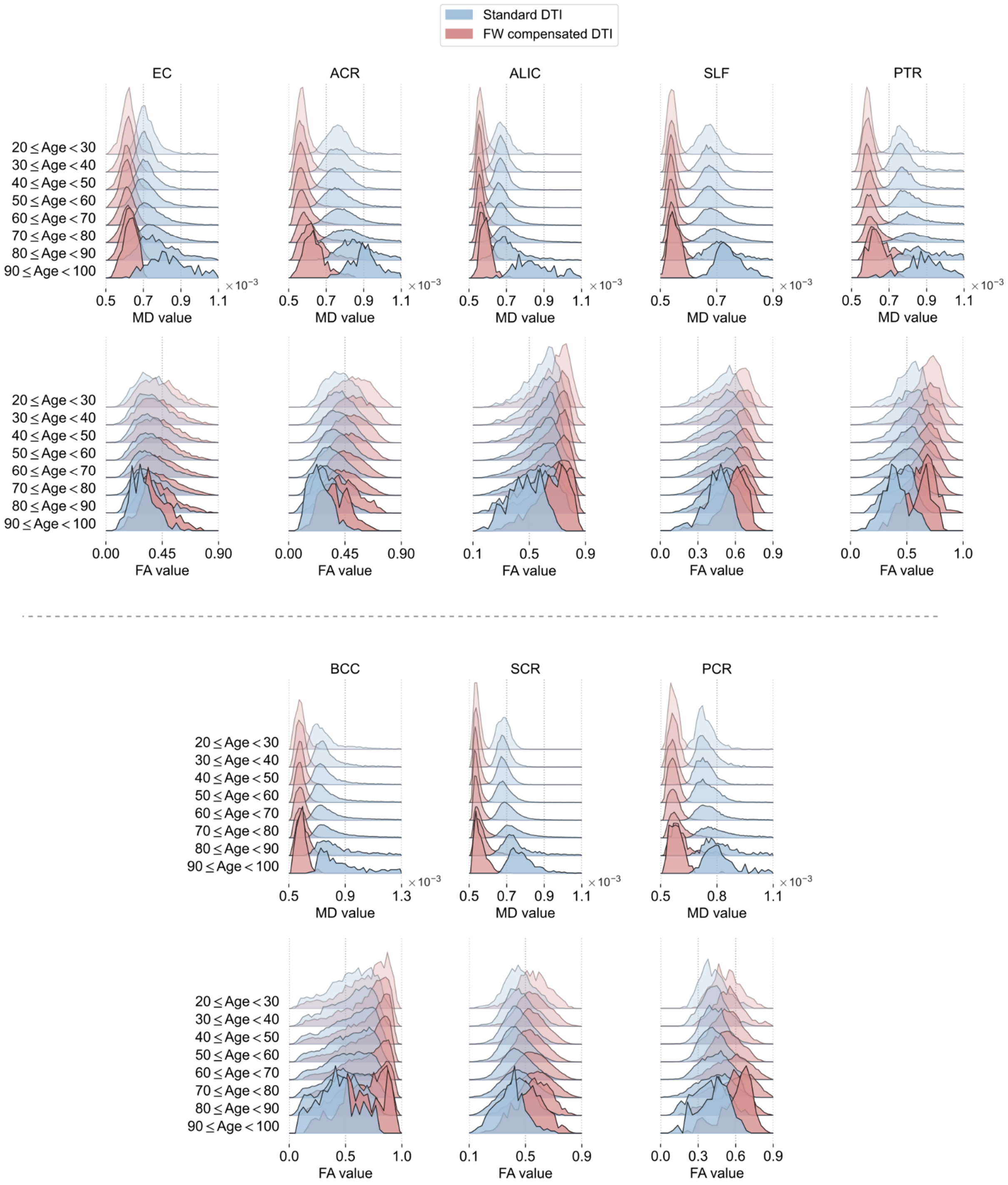


**Supplementary Figure 6.** The histograms presenting the populational variations in MD and FA measures estimated under the standard DTI and FW compensated DTI over the age intervals defined in Table 1. The experiment reports the qualitative MD and FA variations using only cross-sectional samples. The number of histogram bins were determined separately for each age interval using the formula built upon Doane's method (Doane, 1976) (see Methods). The labels use the abbreviations defined in Fig. 1b and Supplementary Fig. 1a,c.

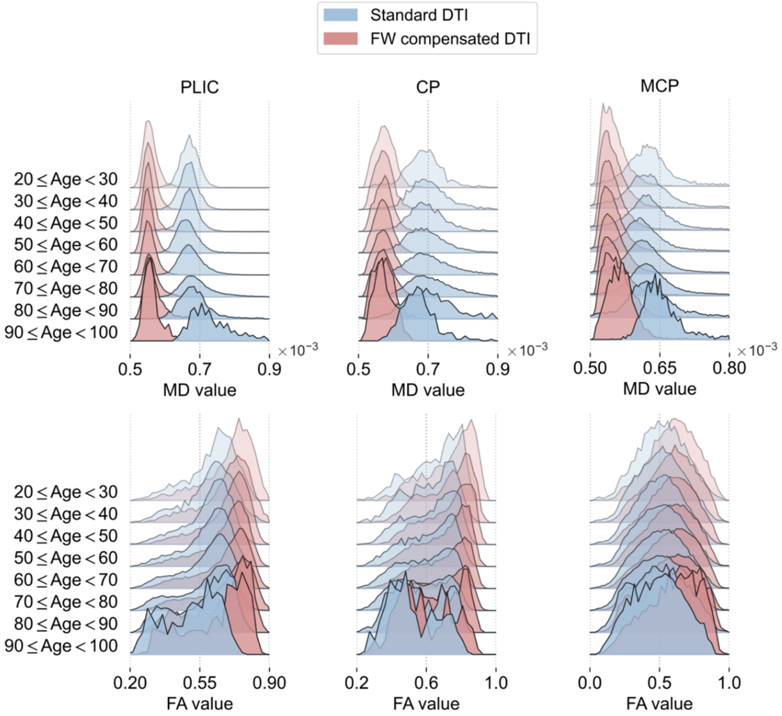


**Supplementary Figure 6. (continued)**

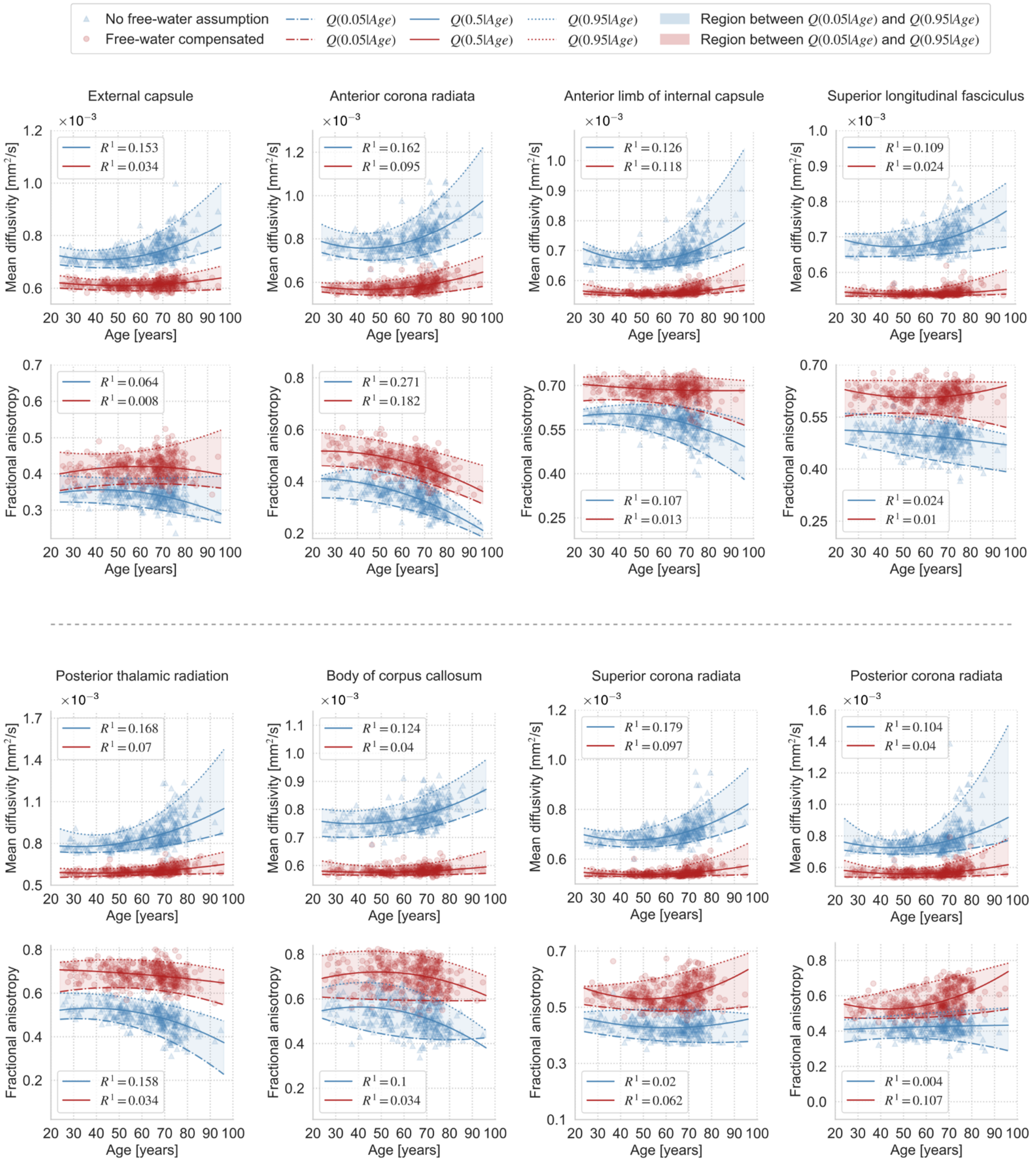


**Supplementary Figure 7.** Estimated MD/FA measures using a standard DTI and with a FW compensation, and their trajectories across the adult lifespan modeled via the QR technique. Each marker represents the median value of the measure calculated over the ROI in the subject's native space. The solid lines show the quantile function $Q(0.5|Age)$, the lower dashed-dotted lines indicate $Q(0.05|Age)$, and the upper dotted lines present $Q(0.95|Age)$, all three computed for a standard DTI (blue lines) and FW compensated DTI (red lines). The regions between $Q(0.05|Age)$ and $Q(0.95|Age)$ were shaded for visualization purposes. The goodness-of-fit $R^1$ at $\tau = 0.5$ was computed separately for a standard DTI and FW compensated DTI measures using the procedure introduced by Koenker & Machado (1999).

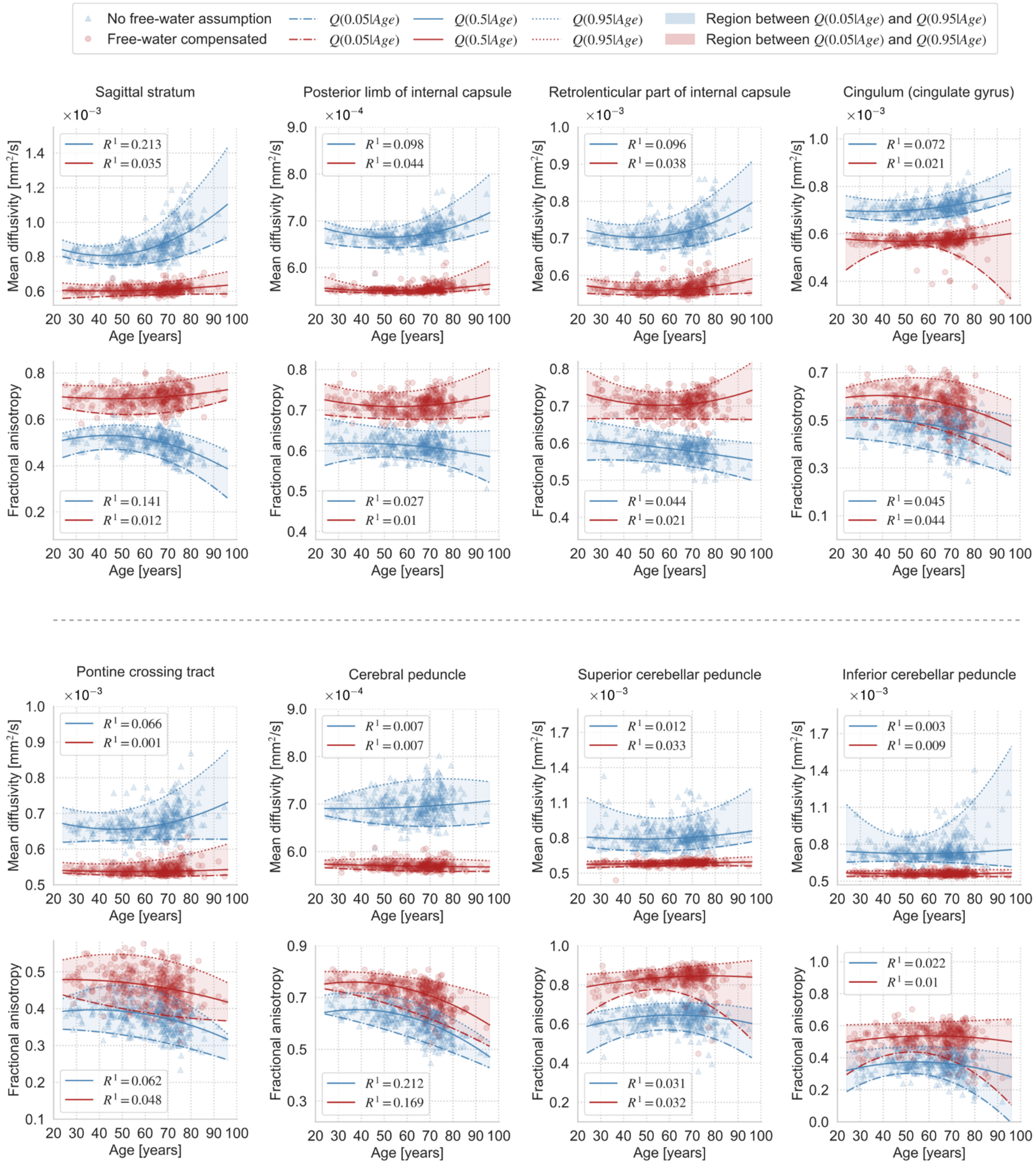


**Supplementary Figure 7. (continued)**

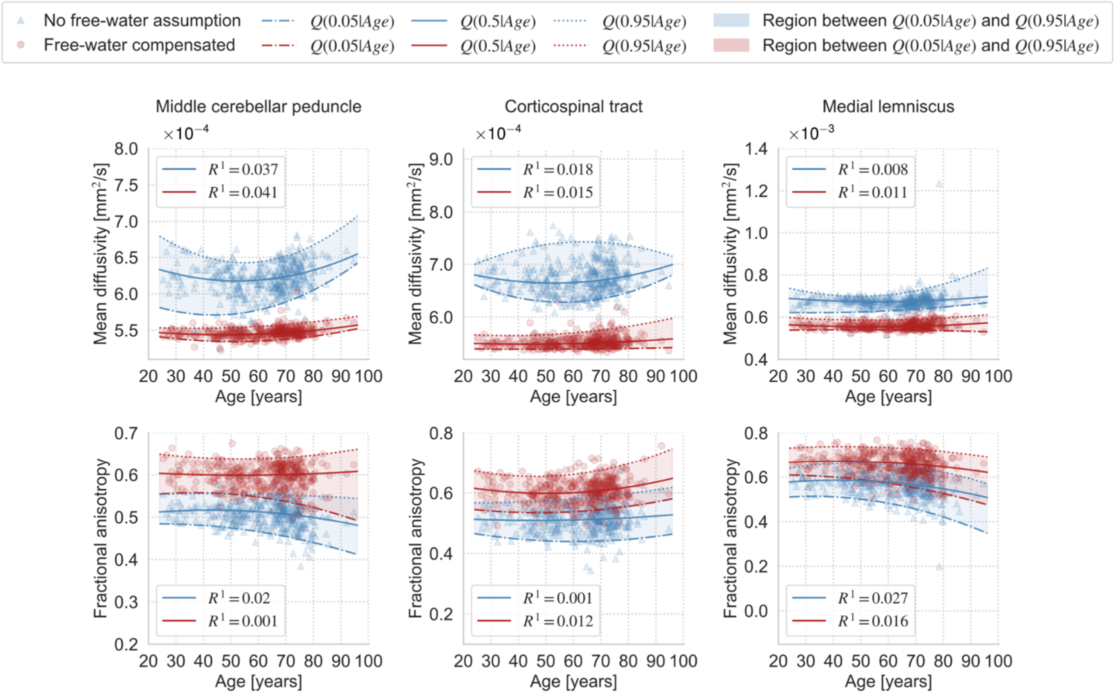


**Supplementary Figure 7. (continued)**

**Supplementary Table 5.** The coefficients of the second-order model fitted to the MD median values from Fig. 4c and Supplementary Fig. 7 using the QR technique under the quantile $\tau = 0.5$, including all regions of interests considered in the study. The columns $\beta_0$, $\beta_1$ and $\beta_2$ represent the constant, linear ($Age$) and quadratic ($Age^2$) terms of the model, respectively. The standard errors of the coefficients are given in the parentheses. The *p*-values refer to the significance of the coefficients: ****p* < .001, ***p* < .01, **p* < .05. If the coefficient is non-significant under the significance level of 0.05 the cell is marked in red and the *p*-value is given in the coefficient superscript. The values over dotted lines refer to the standard MD trajectory (i.e., the estimation was made with no FW assumption), while the bottom ones present the FW compensated MD. The column $R^1\ (\tau)$ includes the goodness-of-fit of the model at $\tau = 0.5$ (Koenker and Machado, 1999). The row names follow the abbreviations defined in Fig. 1b and Supplementary Fig. 1a,c.

| **ROI** | **DTI variant** | **Quantile regression fitting coefficients, *p*-values, standard errors (std. err.) and $R^1\ (\tau)$ at $\tau = 0.5$.** | | | |
|---|---|---|---|---|---|
| | | $\beta_0$ $^{p\text{-value}}$ $\times 10^2$ (std. err. $\times 10^2$) | $\beta_1$ $^{p\text{-value}}$ $\times 10^4$ (std. err. $\times 10^4$) | $\beta_2$ $^{p\text{-value}}$ $\times 10^6$ (std. err. $\times 10^6$) | $R^1\ (\tau)$ |
| WM | standard DTI | 0.077641*** (0.002450) | -0.036626*** (0.008522) | 0.039827*** (0.007385) | 0.172 |
| | FW compensated DTI | 0.058569*** (0.000869) | -0.011404*** (0.003334) | 0.011947*** (0.003032) | 0.094 |
| GCC | standard DTI | 0.108887*** (0.006443) | -0.132227*** (0.022572) | 0.144993*** (0.019333) | 0.224 |
| | FW compensated DTI | 0.065173*** (0.001376) | -0.021939*** (0.005298) | 0.024821*** (0.004780) | 0.124 |
| SCC | standard DTI | 0.077389*** (0.002518) | -0.031862*** (0.008671) | 0.035364*** (0.007182) | 0.146 |
| | FW compensated DTI | 0.061481*** (0.001034) | -0.007140 $^{p = .054}$ (0.003692) | 0.006583* (0.003211) | 0.02 |
| CGh | standard DTI | 0.088538*** (0.007544) | -0.074000* (0.029207) | 0.103632*** (0.027307) | 0.129 |
| | FW compensated DTI | 0.064578*** (0.002758) | -0.008752 $^{p = .355}$ (0.009451) | 0.006683 $^{p = .397}$ (0.007882) | 0.002 |
| EC | standard DTI | 0.078850*** (0.003374) | -0.038370** (0.012679) | 0.045805*** (0.011480) | 0.153 |
| | FW compensated DTI | 0.064670*** (0.001454) | -0.014364** (0.005207) | 0.014118** (0.004462) | 0.034 |
| ACR | standard DTI | 0.090714*** (0.003646) | -0.068958*** (0.014244) | 0.079135*** (0.013399) | 0.162 |
| | FW compensated DTI | 0.062584*** (0.002199) | -0.026520** (0.008366) | 0.029919*** (0.007594) | 0.095 |
| ALIC | standard DTI | 0.081551*** (0.003877) | -0.060284*** (0.014636) | 0.060271*** (0.013300) | 0.126 |
| | FW compensated DTI | 0.059053*** (0.000698) | -0.013726*** (0.002553) | 0.013731*** (0.002259) | 0.118 |

**Supplementary Table 5. (continued)**

| ROI | DTI variant | Quantile regression fitting coefficients, *p*-values, standard errors (std. err.) and $R^1(\tau)$ at $\tau = 0.5$. | | | |
|---|---|---|---|---|---|
| | | $\beta_0$ $^{p\text{-value}}$ $\times 10^2$ (std. err. $\times 10^2$) | $\beta_1$ $^{p\text{-value}}$ $\times 10^4$ (std. err. $\times 10^4$) | $\beta_2$ $^{p\text{-value}}$ $\times 10^6$ (std. err. $\times 10^6$) | $R^1(\tau)$ |
| SLF | standard DTI | 0.075518*** (0.002586) | -0.036360*** (0.008973) | 0.039824*** (0.007650) | 0.109 |
| | FW compensated DTI | 0.055891*** (0.000845) | -0.007797* (0.003054) | 0.007428** (0.002699) | 0.024 |
| PTR | standard DTI | 0.084635*** (0.004614) | -0.043128* (0.018293) | 0.067061*** (0.017473) | 0.168 |
| | FW compensated DTI | 0.061436*** (0.001856) | -0.014463 $^{p = .056}$ (0.007548) | 0.018893* (0.007380) | 0.07 |
| BCC | standard DTI | 0.080645*** (0.002797) | -0.029560** (0.010189) | 0.037831*** (0.008951) | 0.124 |
| | FW compensated DTI | 0.059230*** (0.001180) | -0.006974 $^{p = .100}$ (0.004230) | 0.007550* (0.003681) | 0.04 |
| SCR | standard DTI | 0.078146*** (0.003424) | -0.047917*** (0.012595) | 0.054302*** (0.010974) | 0.179 |
| | FW compensated DTI | 0.057350*** (0.000921) | -0.015176*** (0.003416) | 0.015816*** (0.003090) | 0.097 |
| PCR | standard DTI | 0.088216*** (0.004311) | -0.068836*** (0.015730) | 0.075501*** (0.013944) | 0.104 |
| | FW compensated DTI | 0.063594*** (0.002176) | -0.030090*** (0.008359) | 0.029319*** (0.007654) | 0.04 |
| SS | standard DTI | 0.098854*** (0.005791) | -0.086642*** (0.021527) | 0.102747*** (0.019300) | 0.213 |
| | FW compensated DTI | 0.062000*** (0.001706) | -0.008592 $^{p = .199}$ (0.006683) | 0.010590 $^{p = .085}$ (0.006131) | 0.035 |
| PLIC | standard DTI | 0.073465*** (0.001835) | -0.027405*** (0.006348) | 0.026707*** (0.005469) | 0.098 |
| | FW compensated DTI | 0.056921*** (0.000553) | -0.007175*** (0.001983) | 0.006960*** (0.001712) | 0.044 |
| RPIC | standard DTI | 0.078403*** (0.002568) | -0.035375*** (0.010333) | 0.038164*** (0.009562) | 0.096 |
| | FW compensated DTI | 0.060290*** (0.001614) | -0.016805** (0.005512) | 0.016214*** (0.004626) | 0.038 |

**Supplementary Table 5. (continued)**

| ROI | DTI variant | Quantile regression fitting coefficients, *p*-values, standard errors (std. err.) and $R^1(\tau)$ at $\tau = 0.5$. | | | |
|---|---|---|---|---|---|
| | | $\beta_0$ $^{p\text{-value}}$ $\times 10^2$ (std. err. $\times 10^2$) | $\beta_1$ $^{p\text{-value}}$ $\times 10^4$ (std. err. $\times 10^4$) | $\beta_2$ $^{p\text{-value}}$ $\times 10^6$ (std. err. $\times 10^6$) | $R^1(\tau)$ |
| CGg | standard DTI | 0.072640*** (0.002581) | -0.016772 $^{p=.071}$ (0.009277) | 0.022611** (0.008262) | 0.072 |
| | FW compensated DTI | 0.060266*** (0.002476) | -0.014158 $^{p=.123}$ (0.009158) | 0.014493 $^{p=.077}$ (0.008183) | 0.021 |
| PCT | standard DTI | 0.072281*** (0.001807) | -0.028755*** (0.007049) | 0.030843*** (0.006607) | 0.066 |
| | FW compensated DTI | 0.054589*** (0.001244) | -0.003175 $^{p=.507}$ (0.004778) | 0.002953 $^{p=.499}$ (0.004361) | 0.001 |
| CP | standard DTI | 0.069464*** (0.002471) | -0.002595 $^{p=.784}$ (0.009455) | 0.003981 $^{p=.641}$ (0.008538) | 0.007 |
| | FW compensated DTI | 0.057679*** (0.000554) | -0.001560 $^{p=.429}$ (0.001971) | 0.000654 $^{p=.701}$ (0.001702) | 0.007 |
| SCP | standard DTI | 0.085877*** (0.008146) | -0.028239 $^{p=.323}$ (0.028525) | 0.029523 $^{p=.222}$ (0.024136) | 0.012 |
| | FW compensated DTI | 0.057032*** (0.001362) | 0.001714 $^{p=.721}$ (0.004795) | 0.000963 $^{p=.815}$ (0.004123) | 0.033 |
| ICP | standard DTI | 0.078227*** (0.006164) | -0.021064 $^{p=0.379}$ (0.023891) | 0.019172 $^{p=.385}$ (0.022054) | 0.003 |
| | FW compensated DTI | 0.058436*** (0.001235) | -0.008774* (0.004118) | 0.007310* (0.003355) | 0.009 |
| MCP | standard DTI | 0.067107*** (0.002726) | -0.020408* (0.009307) | 0.019479* (0.007731) | 0.037 |
| | FW compensated DTI | 0.055615*** (0.000590) | -0.005239* (0.002127) | 0.005550** (0.001848) | 0.041 |
| CT | standard DTI | 0.071674*** (0.003018) | -0.019814 $^{p=.060}$ (0.010500) | 0.018757* (0.008676) | 0.018 |
| | FW compensated DTI | 0.055531*** (0.000903) | -0.003024 $^{p=.376}$ (0.003412) | 0.003528 $^{p=.249}$ (0.003054) | 0.015 |
| ML | standard DTI | 0.071926*** (0.002913) | -0.016466 $^{p=.095}$ (0.009829) | 0.014703 $^{p=.071}$ (0.008112) | 0.008 |
| | FW compensated DTI | 0.058453*** (0.001310) | -0.011278* (0.005215) | 0.010444* (0.004820) | 0.011 |

**Supplementary Table 6.** The coefficients of the second-order model fitted to the FA median values from Fig. 4c and Supplementary Fig. 7 using the QR technique under the quantile $\tau = 0.5$, including all regions of interests considered in the study. The columns $\beta_0$, $\beta_1$ and $\beta_2$ represent the constant, linear ($Age$) and quadratic ($Age^2$) terms of the model, respectively. The standard errors of the coefficients are given in the parentheses. The *p*-values refer to the significance of the coefficients: ****p* < .001, ***p* < .01, **p* < .05. If the coefficient is non-significant under the significance level of 0.05 the cell is marked in red and the *p*-value is given in the coefficient superscript. The values over dotted lines refer to the standard FA trajectory (i.e., the estimation was made with no FW assumption), while the bottom ones present the FW compensated FA. The column $R^1$ includes the goodness-of-fit of the model at $\tau = 0.5$ (Koenker & Machado, 1999). The row names follow the abbreviations defined in Fig. 1b and Supplementary Fig. 2a,c.

| ROI | DTI variant | Quantile regression fitting coefficients, *p*-values, standard errors (std. err.) and $R^1(\tau)$ at $\tau = 0.5$. | | | |
|---|---|---|---|---|---|
| | | $\beta_0$ $^{p\text{-value}}$ (std. err.) | $\beta_1$ $^{p\text{-value}} \times 10^2$ (std. err. $\times 10^2$) | $\beta_2$ $^{p\text{-value}} \times 10^4$ (std. err. $\times 10^4$) | $R^1(\tau)$ |
| WM | standard DTI | 0.489697*** (0.018702) | 0.124957 $^{p=.087}$ (0.072938) | -0.173992** (0.066122) | 0.107 |
| | FW compensated DTI | 0.654102*** (0.023021) | -0.115394 $^{p=.196}$ (0.088997) | 0.105635 $^{p=.191}$ (0.080622) | 0.006 |
| GCC | standard DTI | 0.582046*** (0.048547) | 0.519276** (0.182682) | -0.645440*** (0.163256) | 0.174 |
| | FW compensated DTI | 0.901654*** (0.032177) | -0.159433 $^{p=.154}$ (0.111683) | 0.113874 $^{p=.235}$ (0.095793) | 0.012 |
| SCC | standard DTI | 0.752142*** (0.024013) | 0.263692** (0.094748) | -0.309145*** (0.087698) | 0.092 |
| | FW compensated DTI | 0.891167*** (0.017618) | -0.016756 $^{p=.780}$ (0.060046) | 0.017368 $^{p=.728}$ (0.049978) | 0.001 |
| CGh | standard DTI | 0.318384*** (0.033052) | 0.014725 $^{p=.902}$ (0.118964) | -0.198010 $^{p=.055}$ (0.103075) | 0.164 |
| | FW compensated DTI | 0.470647*** (0.048887) | -0.316120 $^{p=.055}$ (0.164355) | 0.245476 $^{p=.070}$ (0.134881) | 0.007 |
| EC | standard DTI | 0.314161*** (0.025894) | 0.196698* (0.088354) | -0.231200** (0.073682) | 0.064 |
| | FW compensated DTI | 0.362583*** (0.026807) | 0.194271* (0.097989) | -0.163939 $^{p=.055}$ (0.085209) | 0.008 |
| ACR | standard DTI | 0.398847*** (0.028230) | 0.130829 $^{p=.200}$ (0.101890) | -0.339688*** (0.089176) | 0.271 |
| | FW compensated DTI | 0.499512*** (0.045281) | 0.153852 $^{p=.315}$ (0.153063) | -0.310211* (0.125901) | 0.182 |
| ALIC | standard DTI | 0.552703*** (0.026132) | 0.267978** (0.095470) | -0.344057*** (0.085538) | 0.107 |
| | FW compensated DTI | 0.723441*** (0.026317) | -0.094154 $^{p=.307}$ (0.092117) | 0.054334 $^{p=.487}$ (0.078091) | 0.013 |

**Supplementary Table 6. (continued)**

| ROI | DTI variant | Quantile regression fitting coefficients, *p*-values, standard errors (std. err.) and $R^1(\tau)$ at $\tau = 0.5$. | | | |
|---|---|---|---|---|---|
| | | $\beta_0$ $^{p\text{-value}}$ (std. err.) | $\beta_1$ $^{p\text{-value}} \times 10^2$ (std. err. $\times 10^2$) | $\beta_2$ $^{p\text{-value}} \times 10^4$ (std. err. $\times 10^4$) | $R^1(\tau)$ |
| SLF | standard DTI | 0.516827*** (0.036319) | -0.008017 $^{p=.951}$ (0.131187) | -0.041903 $^{p=.709}$ (0.112159) | 0.024 |
| | FW compensated DTI | 0.675862*** (0.029371) | -0.249838* (0.102586) | 0.221906* (0.085868) | 0.01 |
| PTR | standard DTI | 0.465260*** (0.028527) | 0.349919** (0.109906) | -0.464518*** (0.099447) | 0.158 |
| | FW compensated DTI | 0.712105*** (0.040685) | -0.002068 $^{p=.988}$ (0.134335) | -0.068224 $^{p=.535}$ (0.109961) | 0.034 |
| BCC | standard DTI | 0.465213*** (0.043723) | 0.485831** (0.150806) | -0.597405*** (0.126661) | 0.1 |
| | FW compensated DTI | 0.609019*** (0.053958) | 0.459086* (0.197799) | -0.471265** (0.174983) | 0.034 |
| SCR | standard DTI | 0.518262*** (0.026854) | -0.298877** (0.103173) | 0.245365** (0.094671) | 0.02 |
| | FW compensated DTI | 0.667070*** (0.034876) | -0.536164*** (0.128866) | 0.522915*** (0.113410) | 0.062 |
| PCR | standard DTI | 0.396867*** (0.038240) | 0.060457 $^{p=.693}$ (0.152960) | -0.023277 $^{p=.873}$ (0.145151) | 0.004 |
| | FW compensated DTI | 0.664293*** (0.055145) | -0.647438** (0.207917) | 0.753470*** (0.188616) | 0.107 |
| SS | standard DTI | 0.428465*** (0.027491) | 0.467474*** (0.110404) | -0.532820*** (0.102684) | 0.141 |
| | FW compensated DTI | 0.723104*** (0.031872) | -0.145429 $^{p=.201}$ (0.113569) | 0.157021 $^{p=.109}$ (0.097638) | 0.012 |
| PLIC | standard DTI | 0.605569*** (0.024539) | 0.070812 $^{p=.424}$ (0.088391) | -0.095175 $^{p=.217}$ (0.076921) | 0.027 |
| | FW compensated DTI | 0.760087*** (0.025750) | -0.184216* (0.089282) | 0.165859* (0.074917) | 0.01 |
| RPIC | standard DTI | 0.621077*** (0.028314) | -0.040503 $^{p=.719}$ (0.112316) | -0.029629 $^{p=.777}$ (0.104369) | 0.044 |
| | FW compensated DTI | 0.788420*** (0.034940) | -0.303192* (0.124066) | 0.265957* (0.107471) | 0.021 |

**Supplementary Table 6. (continued)**

| ROI | DTI variant | Quantile regression fitting coefficients, *p*-values, standard errors (std. err.) and $R^1(\tau)$ at $\tau = 0.5$. | | | |
|---|---|---|---|---|---|
| | | $\beta_0$ ^p-value^ (std. err.) | $\beta_1$ ^p-value^ $\times 10^2$ (std. err. $\times 10^2$) | $\beta_2$ ^p-value^ $\times 10^4$ (std. err. $\times 10^4$) | $R^1(\tau)$ |
| CGg | standard DTI | 0.473763*** (0.070685) | 0.182507 $^{p=.499}$ (0.269728) | -0.278843 $^{p=.263}$ (0.248496) | 0.045 |
| | FW compensated DTI | 0.545547*** (0.067739) | 0.300765 $^{p=.216}$ (0.242744) | -0.388158 $^{p=.067}$ (0.211480) | 0.044 |
| PCT | standard DTI | 0.364030*** (0.026675) | 0.176378 $^{p=.084}$ (0.101809) | -0.234986* (0.092260) | 0.062 |
| | FW compensated DTI | 0.470716*** (0.024522) | 0.064449 $^{p=.490}$ (0.093335) | -0.124647 $^{p=.146}$ (0.085608) | 0.048 |
| CP | standard DTI | 0.573577*** (0.032900) | 0.417484*** (0.117658) | -0.545960*** (0.100162) | 0.212 |
| | FW compensated DTI | 0.697098*** (0.031388) | 0.343168** (0.114727) | -0.469262*** (0.101750) | 0.169 |
| SCP | standard DTI | 0.494364*** (0.041827) | 0.485235** (0.171609) | -0.388267* (0.165148) | 0.031 |
| | FW compensated DTI | 0.727693*** (0.060098) | 0.306550 $^{p=.120}$ (0.196602) | -0.198120 $^{p=.215}$ (0.159632) | 0.032 |
| ICP | standard DTI | 0.205529*** (0.046270) | 0.613002*** (0.171952) | -0.557700*** (0.156497) | 0.022 |
| | FW compensated DTI | 0.429730*** (0.060662) | 0.356255 $^{p=.119}$ (0.228255) | -0.294907 $^{p=.160}$ (0.209259) | 0.01 |
| MCP | standard DTI | 0.493524*** (0.024362) | 0.108237 $^{p=.256}$ (0.095061) | -0.126454 $^{p=.150}$ (0.087689) | 0.02 |
| | FW compensated DTI | 0.613822*** (0.027799) | -0.054483 $^{p=.578}$ (0.097956) | 0.050565 $^{p=.540}$ (0.082508) | 0.001 |
| CT | standard DTI | 0.524498*** (0.043661) | -0.066386 $^{p=.673}$ (0.156986) | 0.072681 $^{p=.598}$ (0.137895) | 0.001 |
| | FW compensated DTI | 0.659435*** (0.052730) | -0.240895 $^{p=.207}$ (0.190637) | 0.238775 $^{p=.148}$ (0.164700) | 0.012 |
| ML | standard DTI | 0.539862*** (0.043115) | 0.221006 $^{p=.153}$ (0.154459) | -0.264757* (0.133964) | 0.027 |
| | FW compensated DTI | 0.644517*** (0.038595) | 0.134478 $^{p=.363}$ (0.147786) | -0.165207 $^{p=.215}$ (0.133059) | 0.016 |
